# Reservoir host immunology and life history shape virulence evolution in zoonotic viruses

**DOI:** 10.1101/2021.10.06.463372

**Authors:** Cara E. Brook, Carly Rozins, Sarah Guth, Mike Boots

## Abstract

The management of future pandemic risk requires a better understanding of the mechanisms that determine the virulence of emerging zoonotic viruses. Meta-analyses suggest that the virulence of emerging zoonoses is correlated with but not completely predictable from reservoir host phylogeny, indicating that specific characteristics of reservoir host immunology and life history may drive the evolution of viral traits responsible for cross-species virulence. In particular, bats host viruses that cause higher case fatality rates upon spillover to humans than those derived from any other mammal, a phenomenon that cannot be explained by phylogenetic distance alone. In order to disentangle the fundamental drivers of these patterns, we develop a nested modeling framework that highlights mechanisms which underpin the evolution of viral traits in reservoir hosts that cause virulence following cross-species emergence. We apply this framework to generate virulence predictions for viral zoonoses derived from diverse mammalian reservoirs, successfully recapturing corresponding virus-induced human mortality rates reported in the literature. Notably, our work offers a mechanistic explanation for the extreme virulence of bat-borne zoonoses and, more generally, demonstrates how key differences in reservoir host longevity, viral tolerance, and constitutive immunity impact the evolution of viral traits that cause virulence following spillover to humans. Our theoretical framework offers a series of testable questions and hypotheses designed to stimulate future work comparing cross-species virulence evolution in zoonotic viruses derived from diverse mammalian hosts.

## Introduction

The devastating impact of the SARS-CoV-2 pandemic highlights the extreme public health outcomes that can result upon cross-species emergence of zoonotic viruses. Estimating the relative threats posed by potential future zoonoses is an important but challenging public health undertaking. In particular, efforts to predict the virulence of emerging viruses can be complicated since chance will always play a role in dictating the initial spillover that precedes selection (1), virulence upon emergence may be maladaptive in novel hosts (2,3), and patterns in available data may be muddled by attainment bias if avirulent infections go underreported (1). Nonetheless, a growing body of recent work highlights clear associations between reservoir and spillover host phylogeny and the virulence of a corresponding cross-species infection (4–7). In many cases, increasing phylogenetic distance between reservoir and spillover hosts is correlated with higher virulence infections (4–6), suggesting that spillover host immune systems may be poorly equipped to tolerate viral traits optimized in more distantly-related reservoirs. Still, the effect of phylogeny on spillover virulence appears to supersede that of simple phylogenetic distance (7,8), indicating that taxon-specific reservoir host immunological and life history traits may be important drivers of cross-species virus virulence. Notably, bats host viruses that cause higher human case fatality rates than zoonoses derived from other mammals and birds, a phenomenon that cannot be explained by phylogenetic distance alone (8). Understanding the mechanisms that select for the evolution of unique viral traits in bats compared to those selected in other mammalian reservoirs should enable us to better predict the virulence of future zoonotic threats.

Although the disproportionate frequency with which the Chiropteran order may source viral zoonoses remains debated (9,10), the extraordinary human pathology induced by many bat-borne zoonoses— including Ebola and Marburg filoviruses, Hendra and Nipah henipaviruses, and SARS, MERS, and SARS-CoV-2 coronaviruses (11)—is not contested. Remarkably, bats demonstrate limited clinical pathology from infection with viruses that cause extreme morbidity and mortality in other hosts (12). Bats avoid pathological outcomes from viral infection via a combination of unique resistance and tolerance mechanisms, which, respectively, limit the viral load accrued during infection (‘resistance’) and reduce the disease consequences of a given viral load (‘tolerance’) (13–16). Viral resistance mechanisms vary across bat species; those described to date include: receptor incompatibilities that limit the extent of infection for certain viruses in certain bats (17–20), constitutive expression of antiviral cytokines in some bat species (21), and enhanced autophagy (22) and heat-shock protein expression (23) in others.

Expansion of anti-viral APOBEC3 genes has also been documented in a few well-studied bat genomes (24,25). While such robust antiviral immunity would result in widespread immunopathology in most mammals, bats—as the only mammals capable of powered flight—have evolved numerous unique mechanisms of mitigating inflammation incurred during the intensive physiological process of flying (26–28). These anti-inflammatory adaptations include loss of PYHIN (29–31) and downregulation of NLRP3 (32) inflammasome-forming gene families, loss of pro-inflammatory genes in the NF-*κB* pathway (24), dampened interferon activation in the STING pathway (33), and diminished caspase-1 inflammatory signaling (34). In addition to facilitating flight, this resilience to inflammation has yielded the apparent by-products of extraordinarily long bat lifespans (35) and tolerance of the immunopathology that typically results from viral infection (11). Moreover, recent work demonstrates how high virus growth rates easily tolerated in constitutively antiviral bat cells cause significant pathology in cells lacking these unique antiviral defenses (36). The extent to which inflammatory tolerance may modulate the evolution of the viruses that bats host, however, remains largely unexplored.

Modern theory on the evolution of virulence typically assumes, either explicitly or implicitly, that high pathogen growth rates should both enhance between-host transmission and elevate infection-induced morbidity or mortality, resulting in a trade-off between virulence and transmission (37–39). Theory further suggests that because viral ‘tolerance’ mitigates virulence without reducing viral load, most host strategies of tolerance should select for higher growth rate pathogens that achieve gains in between-host transmission without causing damage to the original host (39–41). The widely touted viral tolerance of bats (11,16,42–44) should therefore be expected to support the evolution of enhanced viral growth rates, which—though avirulent to bats—may cause significant pathology upon spillover to hosts lacking unique features of bat immunology and physiology. Beyond tolerance, other life history characteristics unique to diverse reservoir hosts should also impact the evolution of traits in the viruses they host—with important consequences for cross-species virulence following spillover. However, to date, we lack a specific theory that examines the relative impact of reservoir host life history on spillover virulence. Here, we explore the extent to which the immunological and life-history traits of mammalian reservoirs can explain variation in the virulence of zoonotic viruses emerging into human hosts.

## Results

### General trends in the evolution of high growth rate viruses

To elucidate how immunological and life-history traits of mammalian hosts combine to drive zoonotic virus virulence, we adopted a nested modeling approach (45), embedding a simple within-host model of viral and leukocyte dynamics within an epidemiological, population-level framework (Fig 1). We examined how the life history traits of a primary reservoir drive the evolution of viral traits likely to cause pathology in a secondary, spillover host—chiefly, a human. Using our nested model in an adaptive dynamics framework (46), we first expressed the conditions—called the ‘invasion fitness’—which permit invasion of an evolutionarily ‘fitter’, mutant virus into a reservoir host system in within-host terms. From this, we derived an expression for 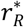, the optimal within-host growth rate of a persistent virus evolved at equilibrium prevalence in its reservoir host; our modeling framework allowed us to express 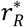 as a function of within-host traits that we might expect to vary across mammalian reservoirs with divergent life histories. We deduced that 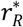 can, consequently, also be expected to vary as a result of these life history differences, which modulate the optimization process by which a virus maximizes gains in between-host transmission 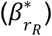 while mitigating virulence incurred on its reservoir (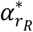; Fig 1). From this framework, we next derived an expression for *α*_*S*_, the virulence incurred by a reservoir-optimized virus immediately following spillover to a novel host. We expressed *α*_*S*_ as a function of the reservoir-optimized virus growth rate 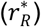, combined with spillover host tolerance of virus pathology (*T*_*vS*_), which we modeled as proportional to the phylogenetic distance between reservoir and spillover host (Fig 1).

**Fig 1.**
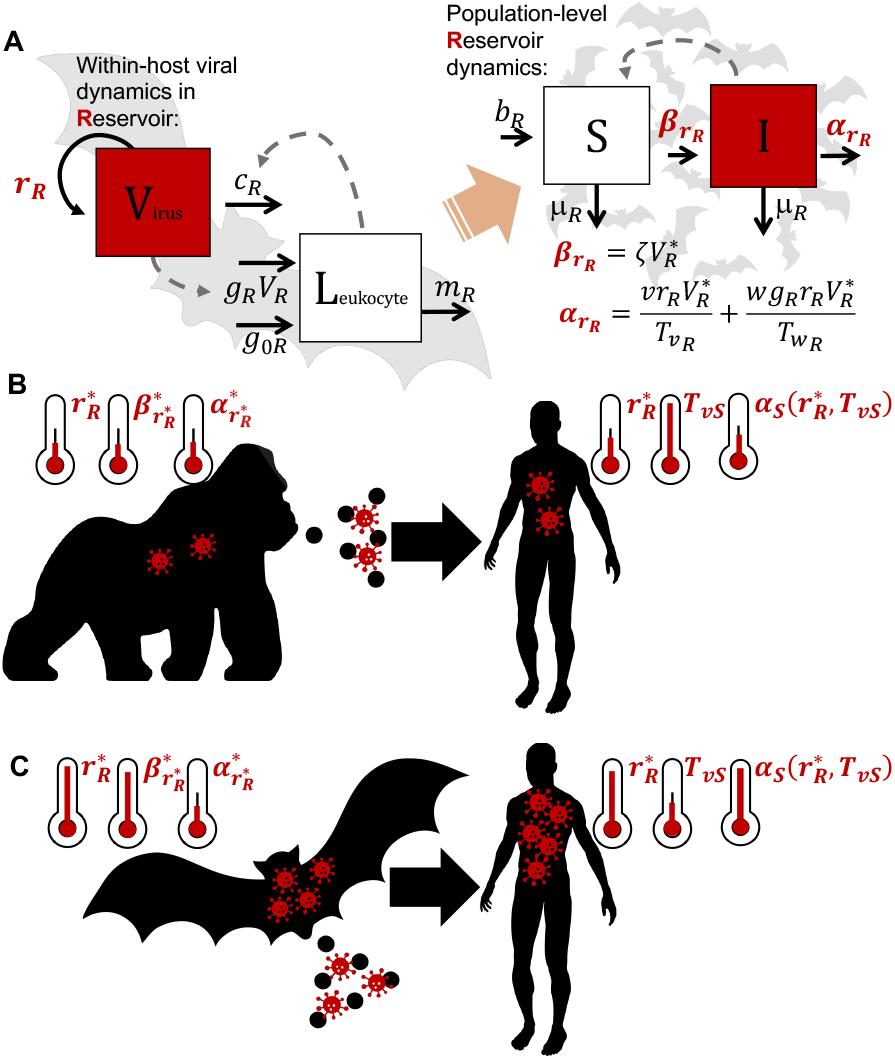
Conceptual mechanistic framework to predict zoonotic virus virulence from reservoir immunology and life history traits. (A) A within-host predator-prey-like model of leukocyte-virus dynamics is embedded in a population-level transmission model for reservoir hosts. Between-host rates of transmission 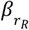 and virulence 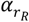 are expressed as functions of within-host dynamics to derive optimal virus growth rates in the reservoir 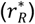. See S1 File and Table 1 for parameter definitions and values. (B) Virus growth rates 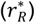 evolve to maximize gains in between-host transmission 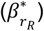 over virulence incurred on the reservoir host 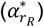 where both transmission and virulence are expressed as functions of *r*_*R*_. Viruses evolved in taxa with immunological and physiological environments similar to humans (i.e. non-human primates) should optimize at relatively low growth rates 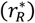 that yield minimal pathology for the reservoir host 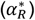 and are, correspondingly, likely to cause only minimal pathology in the spillover human host (*α*_*S*_). In our model, the virulence of a virus in its spillover host (*α*_*S*_) is expressed as a function of both the reservoir host-evolved virus growth rate 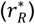 and the spillover host tolerance of virus pathology (*T*_*vS*_), which we model as proportional to phylogenetic distance between reservoir and spillover host. (C) As a result of unique bat virus tolerance, viruses evolved in bat reservoir hosts may optimize at high 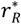 values that maximize bat-to-bat transmission 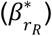 but cause only minimal pathology in the bat host 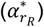. Such viruses are likely to generate extreme virulence upon spillover (*α*_*S*_) to secondary hosts, including humans, that lack bat life history traits. The virulence of a virus in its spillover host is amplified (*α*_*S*_) in cases where large phylogenetic distance between reservoir and spillover host results in minimal spillover host tolerance of virus pathology (*T*_*vS*_).

Our nested modeling approach first followed Alizon and van Baalen (2005) (45) in derivation of an expression for 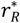, the within-host virus growth rate optimized based on endemic circulation in a reservoir host. The equation for 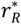 can be captured as follows:

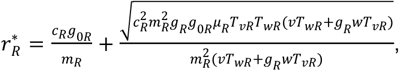

where *μ*_*R*_ signifies the natural mortality rate of the reservoir host, and all other parameters represent within-host viral and immune dynamics in a persistently infected reservoir. Thus, *c*_*R*_ corresponds to the rate of virus clearance by leukocytes, *g*_0*R*_ is the magnitude of constitutive immunity, *m*_*R*_ is the natural leukocyte mortality rate, and *g*_*R*_ is the rate of leukocyte activation following infection, all in the reservoir host. The parameters *v* and *w* correspond, respectively, to the intrinsic virulence of the virus and its propensity to elicit a damaging inflammatory response from its host’s immune system—while *T*_*vR*_ and *T*_*wR*_ respectively represent reservoir host tolerance to direct virus pathology and to immunopathology. From the above expression, we explored a range of optimal within-host virus growth rates 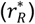 for viruses evolved in reservoir hosts with diverse cellular and immunological parameters (Fig 2; Table 1; S1-S2 Figs). We subsequently calculated the corresponding transmission 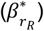 and virulence 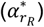 incurred by viruses evolved to 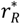 in their reservoir hosts, then, following Gilchrist and Sasaki (2002) (47), modeled the nascent spillover of a reservoir-evolved virus as acute infection in a secondary host. In this spillover infection, we assumed that the virus retains its reservoir-optimized growth rate 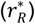 while replicating in the physiological and immunological environment of its novel spillover host. We express this spillover virulence (*α*_*S*_) as:

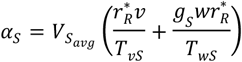

where 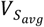 corresponds to the average viral load experienced across the timecourse of an acute spillover host infection, and *g*_*S*_, *T*_*vS*_, and *T*_*wS*_ represent the spillover host analogues of previously-described within-host parameters in the reservoir host equations (Fig 2; Table 1; S1-S2 Figs). We present all main text results under assumptions by which tolerance manifests as a *constant* reduction of either direct virus-induced pathology (*T*_*v*_) or immunopathology (*T*_*w*_). See S1 File for comparable results under assumptions of *complete* tolerance, whereby tolerance completely eliminates virus pathology and immunopathology up to a threshold value, beyond which pathology scales proportionally with virus and immune cell growth.

**Fig 2.**
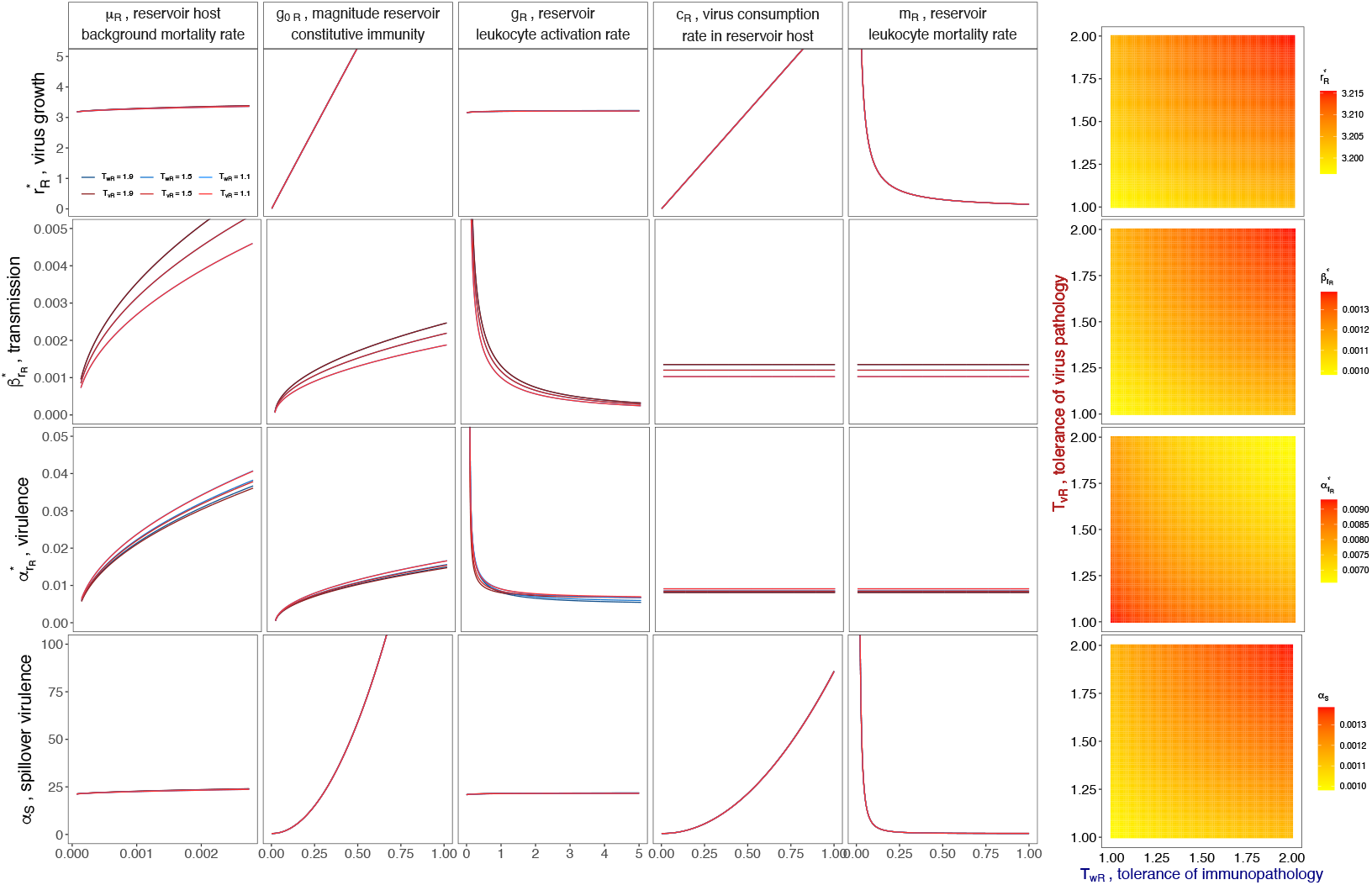
Optimal virus growth rates—and subsequent spillover virulence—vary across reservoir host immunological and life history parameters. Rows (top-down) indicate the evolutionarily optimal within-host virus growth rate 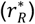 and the corresponding between-host transmission rate 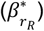, and virus-induced mortality rate 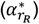 for a reservoir host infected with a virus at 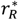. The bottom row then demonstrates the resulting virulence (*α*_*S*_) of a reservoir-optimized virus evolved to 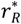 upon nascent spillover to a novel, secondary host. Columns demonstrate the dependency of these outcomes on variable within-host parameters in the reservoir host: background mortality (*μ*_*R*_), magnitude of constitutive immunity (*g*_0*R*_), rate of leukocyte activation upon viral contact (*g*_*R*_), rate of virus consumption by leukocytes (*c*_*R*_), and leukocyte mortality rate (*m*_*R*_). Darker colored lines depict outcomes at higher reservoir host tolerance of direct virus pathology (*T*_*vR*_,red) or immunopathology (*T*_*wR*_,blue), assuming no tolerance of the opposing type. Heat maps demonstrate how *T*_*vR*_ and *T*_*wR*_ interact to produce each outcome. Parameter values are reported in Table 1. Figure assumes tolerance in the ‘constant’ form. See S2 Fig for ‘complete’ tolerance assumptions and S3 Fig for changes in *α*_*S*_ across a variable range of parameter values for *spillover* host tolerance of direct virus pathology, *T*_*vS*_, and immunopathology, *T*_*wS*_.

**Table 1.**
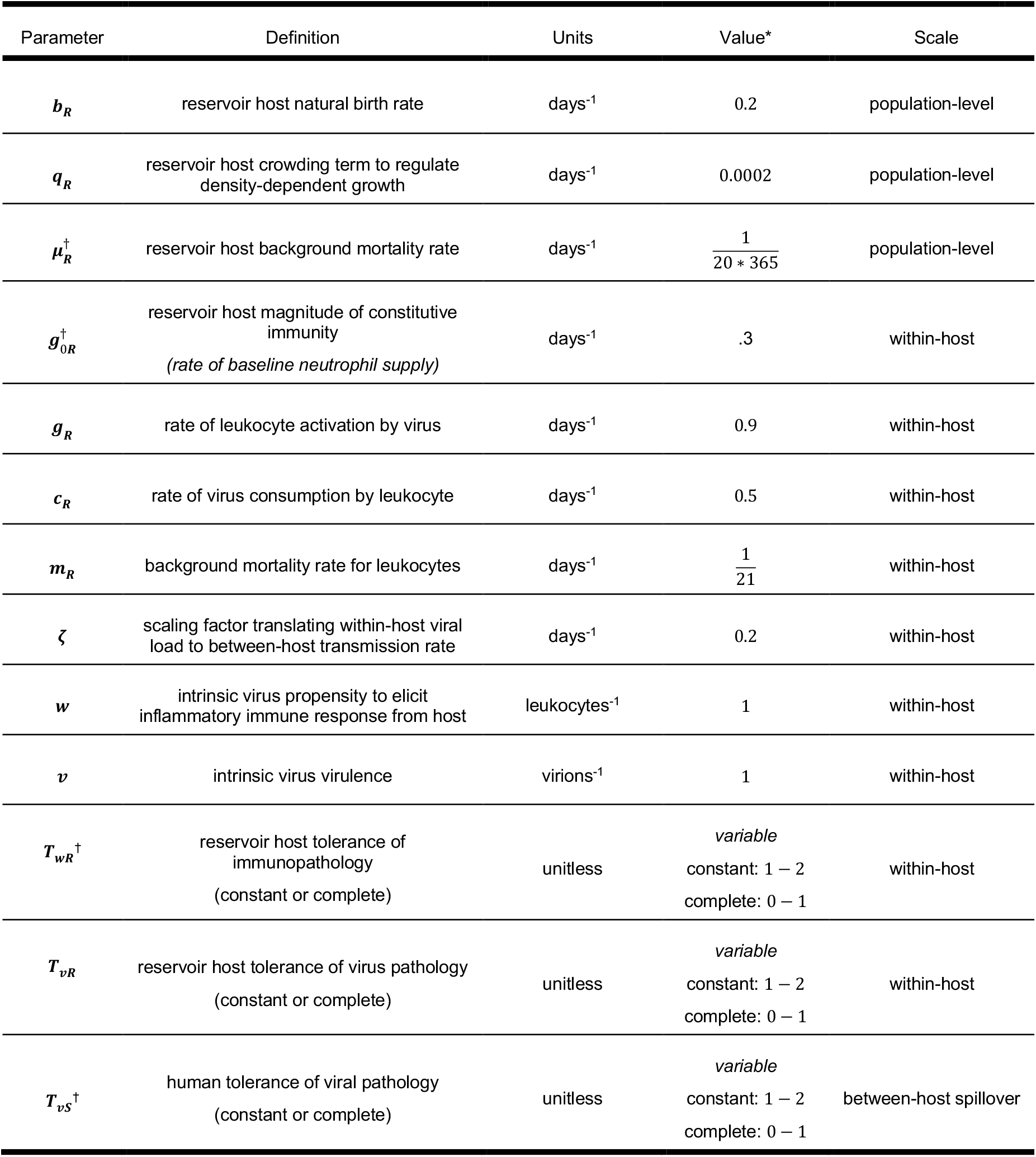
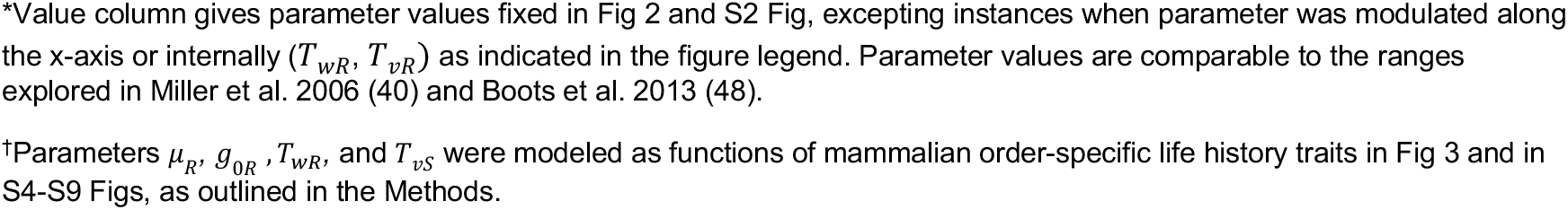
Default parameter values for population-level and within-host models.

| Parameter | Definition | Units | Value* | Scale |
| --- | --- | --- | --- | --- |
| $b_R$ | reservoir host natural birth rate | days<sup>-1</sup> | 0.2 | population-level |
| $q_R$ | reservoir host crowding term to regulate density-dependent growth | days<sup>-1</sup> | 0.0002 | population-level |
| $\mu_R^\dagger$ | reservoir host background mortality rate | days<sup>-1</sup> | $\frac{1}{20 * 365}$ | population-level |
| $g_{0R}^\dagger$ | reservoir host magnitude of constitutive immunity<br>(rate of baseline neutrophil supply) | days<sup>-1</sup> | .3 | within-host |
| $g_R$ | rate of leukocyte activation by virus | days<sup>-1</sup> | 0.9 | within-host |
| $c_R$ | rate of virus consumption by leukocyte | days<sup>-1</sup> | 0.5 | within-host |
| $m_R$ | background mortality rate for leukocytes | days<sup>-1</sup> | $\frac{1}{21}$ | within-host |
| $\zeta$ | scaling factor translating within-host viral load to between-host transmission rate | days<sup>-1</sup> | 0.2 | within-host |
| $w$ | intrinsic virus propensity to elicit inflammatory immune response from host | leukocytes<sup>-1</sup> | 1 | within-host |
| $v$ | intrinsic virus virulence | virions<sup>-1</sup> | 1 | within-host |
| $T_{wR}^\dagger$ | reservoir host tolerance of immunopathology<br>(constant or complete) | unitless | *variable*<br>constant: 1 – 2<br>complete: 0 – 1 | within-host |
| $T_{vR}$ | reservoir host tolerance of virus pathology<br>(constant or complete) | unitless | *variable*<br>constant: 1 – 2<br>complete: 0 – 1 | within-host |
| $T_{vS}^\dagger$ | human tolerance of viral pathology<br>(constant or complete) | unitless | *variable*<br>constant: 1 – 2<br>complete: 0 – 1 | between-host spillover |

\*Value column gives parameter values fixed in Fig 2 and S2 Fig, excepting instances when parameter was modulated along the x-axis or internally ( $T_{wR}$ , $T_{vR}$ ) as indicated in the figure legend. Parameter values are comparable to the ranges explored in Miller et al. 2006 (40) and Boots et al. 2013 (48).
†Parameters $\mu_R$ , $g_{0R}$ , $T_{wR}$ , and $T_{vS}$ were modeled as functions of mammalian order-specific life history traits in Fig 3 and in S4-S9 Figs, as outlined in the Methods.

Our analyses highlight several critical drivers of virus evolution likely to generate significant pathology following spillover to a secondary host (Fig 2): higher within-host virus growth rates 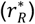 are selected in reservoir hosts with higher background mortality (*μ*_*R*_), elevated constitutive immune responses (*g*_0*R*_), more rapid leukocyte activation upon infection (*g*_*R*_), and more rapid virus clearance by the host immune system (*c*_*R*_). Additionally, higher 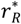 viruses are selected in hosts exhibiting lower leukocyte mortality rates (*m*_*R*_), resulting in longer-lived immune cells. Critically, higher reservoir host tolerance of both virus-induced pathology (*T*_*vR*_) and immunopathology (*T*_*wR*_) also select for higher 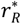. In keeping with trade-off theory, changes in the majority of within-host parameters drive corresponding increases in 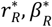, and 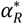, such that viruses evolved to high growth rates experience high transmission within the reservoir host population—but also generate high virus-induced mortality (Fig 2). By definition, the two modeled mechanisms of tolerance (*T*_*vR*_ and *T*_*wR*_) decouple the relationship between transmission and virulence, permitting evolution of high 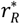 viruses that achieve gains in between-host transmission 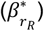, while simultaneously incurring minimal virulence 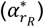 on reservoir hosts. By extension, we demonstrate that viruses evolving high optimal 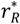 values in reservoir hosts incur substantial pathology upon spillover to secondary hosts (*α*_*S*_). Intriguingly, the virulence that a virus incurs on its spillover host (*α*_*S*_) accelerates substantially faster than that incurred on its reservoir host 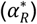 at higher values for certain parameters, chiefly, *g*_0*R*_, *c*_*R*_ as well as, most critically, *T*_*vR*_ and *T*_*wR*_. This underscores the important capacity of these within-host traits to drive cross-species virulence in emerging viruses.

### Order-specific estimates for optimal virus growth rates evolved in reservoir hosts

We next applied our model to make broad predictions of the evolution of optimal virus growth rates 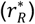 across diverse mammalian reservoir orders, based on order-specific variation in three key parameters from our nested model: the reservoir host background mortality rate (*μ*_*R*_), the reservoir host tolerance of immunopathology (*T*_*wR*_), and the reservoir host magnitude of constitutive immunity (*g*_0*R*_). Within-host immunological data needed to quantify these parameters is lacking for most taxa; thus, we used regression analyses to summarize these terms across mammalian orders from publicly available life history data. In particular, we used well-described allometric relationships between mammalian body mass and basal metabolic rate (BMR) with lifespan and immune cell concentrations (49–53)—to proxy *μ*_*R*_, *T*_*wR*_, and *g*_0*R*_ across mammalian orders (Fig 3A-C; Table 1; S4-S5 Figs). From here, we used our nested modeling framework to predict optimal virus growth rates 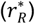 across diverse mammalian reservoirs (Fig 3D). Then, we estimated 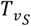, the spillover host tolerance of direct virus pathology, as proportional to the time to most recent common ancestor (MRCA) between the human primate order and each mammalian reservoir order, now focusing our analysis on zoonotic spillover (Fig 3E-F; Table 1). Finally, we combined estimates for 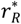 and 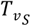 to generate a prediction for *α*_*S*_, the virulence of a reservoir-optimized virus in a human spillover host, which we can compare against case fatality rates for mammalian zoonoses reported in the literature (4,8) (Fig 3G; S5-S9 Figs).

**Fig 3.**
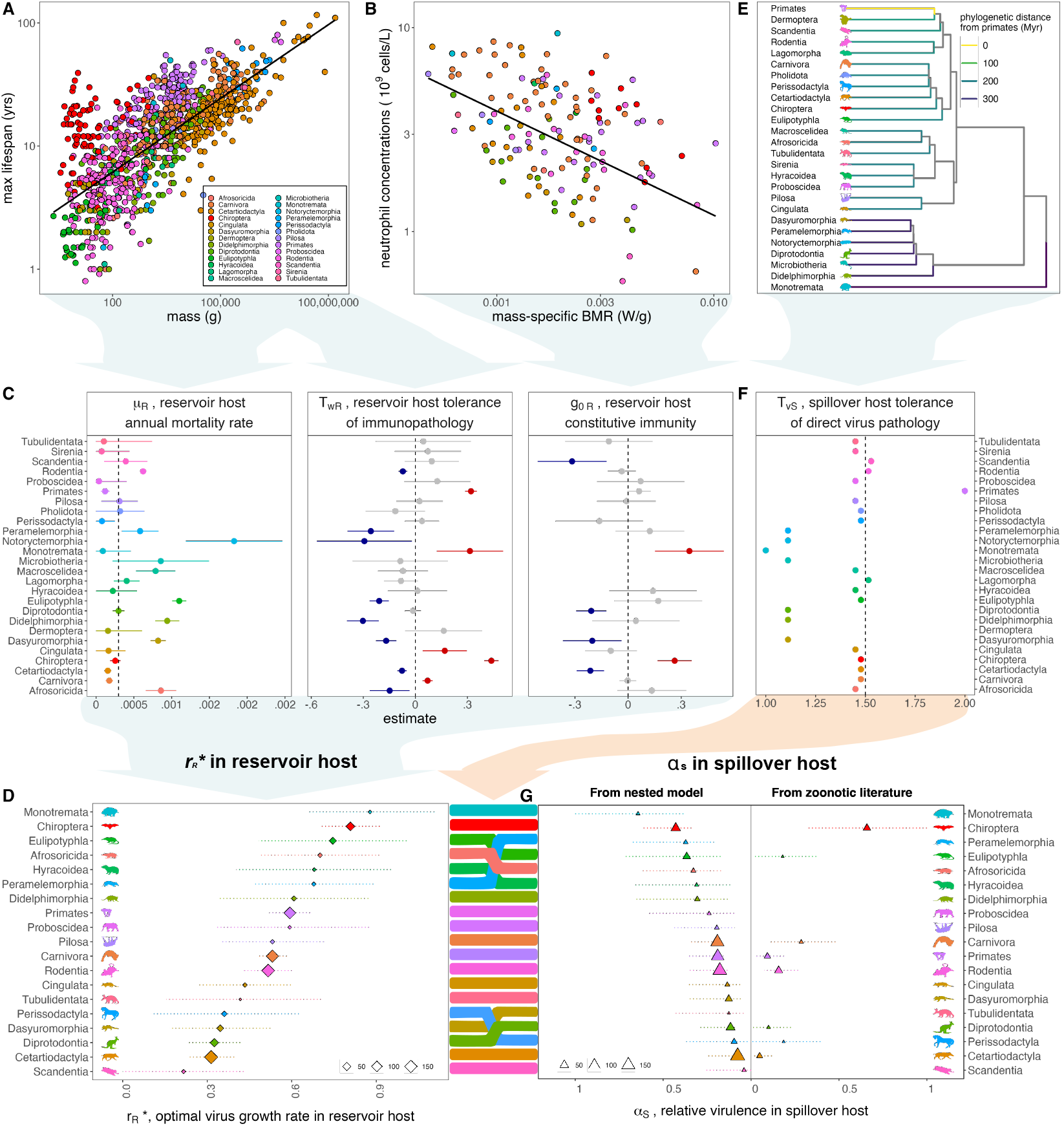
Reservoir host life history traits predict evolution of zoonotic virus virulence. (**A**) Variation in log_10_ maximum lifespan (y-axis, in years) with log_10_ adult body mass (x-axis, in grams) across mammals, with data derived from Jones et al. 2009 (54) and Healy et al. 2014 (55). Points are colored by mammalian order, corresponding to legend. Black line depicts predictions of mammalian lifespan per body mass, summarized from fitted model (but excluding random effect of mammalian order), presented in S2 Table. (**B**) Baseline neutrophil concentrations (y-axis, in 10^9^ cells/L) per mass-specific metabolic rate (x-axis, in W/g) across mammals, with data from Jones et al. 2009 (54) and Healy et al. 2014 (55) combined with neutrophil concentrations from Species360 (53). Black line projects neutrophil concentration per mass-specific metabolic rate (excluding random effects of mammalian order), simplified from fitted model presented in S2 Table. (**C**) Order-level parameters for nested modeling framework were derived from fitting of linear models and linear mixed models visualized here and presented in S4 Fig and S2 Table to data from (A) and (B). Average annual mortality rate (*μ*_*R*_) was predicted from a linear regression of species-level annual mortality (the inverse of maximum lifespan), as described by a predictor variable of host order; tolerance of immunopathology 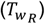 was derived from the scaled effect of host order on the linear mixed effects regression of log_10_ maximum lifespan (in years) by log_10_ mass (in grams), incorporating a random effect of order. The magnitude of constitutive immunity 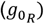 was derived from the scaled effect of order on the regression of log_10_ neutrophil concentration per log_10_ body mass (in grams), combined with BMR (in W) (S4 Fig; S1-S2 Tables). Panels shown here give numerical estimates for *μ*_*R*_ and order-level effects from fitted models that were scaled to numerical values for 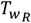 and 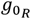,as presented in S4 Fig (S1 Table). Red and blue colors correspond to, respectively, significantly positive or negative order-level partial effects from these regressions. (**D**) Reservoir-host estimates for *μ*_*R*_, *T*_*wR*_, and *g*_0*R*_ were combined in our modeling framework to generate a prediction of optimal growth rate for a virus evolved in a host of each mammalian order 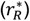. Here, point size corresponds to the average number of species-level datapoints used to generate each of the three variable parameters impacting 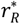, as indicated in legend. (**E**) Phylogenetic distance from Primates (in millions of years, indicated by color) on a timescaled phylogeny, using data from TimeTree (56). (**F**) An order-level estimate for the nested model parameter, *T*_*vS*_, the spillover human host tolerance of pathology induced by a virus evolved in a different reservoir order, was estimated as the scaled inverse of the phylogenetic distance shown in (E) (S4 Fig; S1 Table). (**G**) Reservoir-host predictions of optimal virus growth rates 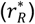 from (D) were combined with human spillover host estimates of tolerance for direct virus pathology (*T*_*vS*_) from (F) in our nested modeling framework to generate a prediction of the relative spillover virulence (*α*_*S*_) of a virus evolved in a given reservoir host order immediately following spillover into a secondary, human host. Here, the left panel visualizes predictions from our nested modeling framework, using order-specific parameters for *μ*_*R*_, *T*_*wR*_, *g*_0*R*_, and *T*_*vS*_ (S1 Table). The right panel depicts relative human *α*_*S*_ estimates derived from case fatality rates and infection duration reported in the zoonotic literature (8). For the left panel, point size corresponds to the average number of species-level datapoints used to generate each of the four variable parameters impacting *α*_*S*_. For the right panel, point size indicates the total number of independent host-virus associations from which virulence estimates were determined. In (C), (D), and (G), 95% confidence intervals were computed by standard error; in (G) for the left panel, these reflect the upper and lower confidence intervals of the optimal virus growth rate in (D). See S1 Table for order-level values for 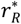, *μ*_*R*_, *T*_*wR*_, *g*_0*R*_, and *T*_*vS*_ and Table 1 for all other default parameters involved in calculation of *α*_*S*_. Sensitivity analyses for zoonotic predictions are summarized in S5-S9 Figs and S3 Table.

We first fit a simple linear regression to the response variable of the inverse of maximum lifespan across diverse mammalian hosts with a single predictor of reservoir host order (Fig 3A; S2 Table). From this simple model, we easily made projections that summarized *μ*_*R*_, the reservoir host annual mortality rate, to an average value for each of 26 mammalian orders. We expressed *μ*_*R*_ in units of days^-1^ on a timescale most relevant to viral dynamics; as a result, even multi-year differences in maximum longevity did little to drive differences in *μ*_*R*_ across mammalian orders. Nonetheless, eight orders (Afrosoricida, Dasyuromorphia, Didelphimorphia, Eulipotyphla, Macroscelidea, Notoryctemorphia, Peramelemorphia, Rodentia) demonstrated significantly elevated annual mortality rates using the median order mortality rate (Diprotodontia) as a reference. By contrast, Carnivora, Cetartiodactyla, Primates, and Perissodactyla demonstrated significantly reduced annual mortality rates as compared to the same reference (Fig 3C; S1-S2 Tables).

Drawing on well-described allometric relationships between mass and lifespan (49) and more recent literature that links longevity with resilience to inflammation (11,44,57–61), we next sought to characterize variation in *T*_*wR*_, reservoir host tolerance of immunopathology, across mammalian orders. Smaller-bodied organisms are hypothesized to be shorter-lived due to higher metabolic rates, more rapid energy expenditure, and faster accumulation of oxidative damage, which can manifest as inflammation (62). We hypothesized, then, that organisms that are longer-lived than predicted for their body size might be more resilient to inflammatory stressors—and, therefore, by extension, more tolerant of immunopathology. Building on this hypothesis, we scaled *T*_*wR*_ after order-level deviations in lifespan as predicted by body mass. To this end, we fit a linear mixed effect regression with a random effect of host order to the log_10_ relationship of lifespan (in years) as predicted by body size (in grams), again spanning data representing 26 diverse mammalian orders. From this, we identified five orders (Carnivora, Chiroptera, Cingulata, Monotremata, and Primates) with significantly longer lifespans than predicted by body size—which scaled to enhanced estimates for *T*_*wR*_. By contrast, we identified eight orders (Afrosoricida, Cetartiodactyla, Dasyuromorphia, Didelphimorphia, Eulipotyphla, Notoryctemorphia, Peramelemorphia, and Rodentia) with significantly shorter lifespans than predicted by body size, which scaled to lower value estimates for *T*_*wR*_ (Fig 3A,C; S4 Fig; S1-S2 Tables). Because the same data on host lifespan were included in both estimation of *μ*_*R*_ and *T*_*wR*_, order-level estimates for *T*_*wR*_ largely mirrored those for *μ*_*R*_—such that longer-lived orders were modeled with low values for annual mortality rate and high values for tolerance to immunopathology (and vice versa). However, because mass was not factored into estimation of *μ*_*R*_, these parameters could diverge in select cases: for example, we estimated mid-range mortality rates for order Chiroptera, as compared with other mammals, but very high values for *T*_*wR*_ becase Chiropteran lifespans—though not remarkably long at face value—far exceed those predicted by body size. By contrast, we estimated low values for both *μ*_*R*_ and *T*_*wR*_ for order Cetartiodactyla, as hosts in this order are long-lived but less long-lived than predicted for body size.

Finally, we fit another linear mixed effect regression with a random effect of host order to the log_10_ relationship of baseline circulating neutrophil concentration (in 10^9^ cells/L) as predicted by mass (in grams) and basal metabolic rate (W); these data spanned only 19 mammalian orders (Fig 3B). From this, we identified a significant positive association between the orders Chiroptera and Monotremata and the response variable of baseline neutrophil concentration, indicating that species in these orders may have more enhanced constitutive immune responses than predicted by mass-specific BMR (Fig 3C). The orders Cetartiodactyla, Dasyuromorphia, Diprotodontia, and Scandentia, by contrast, showed significant negative associations, representing lower baseline neutrophil concentrations than predicted per mass-specific BMR. We scaled these order-level effects to correspondingly high and low estimates for *g*_0*R*_, the within-host parameter representing the magnitude of reservoir-host constitutive immunity in our nested modeling framework (Fig 3B,C; S4 Fig; S1-S2 Tables).

From life history-derived estimates for *μ*_*R*_, *T*_*wR*_, and *g*_0*R*_, we next generated a prediction of the optimal growth rate of a virus evolved in a reservoir host 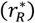 for each of 19 distinct mammalian orders for which we possessed the full suite of proxy data for all three variable within-host parameters (Fig 3D). In keeping with results from our general model (Fig 2), we predicted the evolution of high 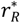 viruses from reservoir orders exhibiting high *μ*_*R*_, *T*_*wR*_, and/or *g*_0*R*_.Because our nested model expressed parameters in timesteps most relevant to viral dynamics (days), differences in mortality rates across mammalian orders—though substantial on multi-year timescales of the host—had limited influence on downstream predictions of differences in the evolution of virus growth rates. This result echoed prior observations from Fig 2, which showed reduced sensitivity of 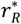 to realistic variation in the magnitude of *μ*_*R*_, as compared with other parameters. As a result, we ultimately recovered the highest predicted optimal growth rates for viruses evolved in the orders Chiroptera and Monotremata, which exhibited both long lifespans per body size (corresponding to high estimates for *T*_*wR*_) and high baseline neutrophil concentrations (corresponding to high estimates for *g*_0*R*_). Data were particularly sparse, however, for order Monotremata, for which complete records were only available for two species (the platypus and the short-beaked echidna). As a result, predictions for this order should be interpreted with caution. More data, particularly for baseline neutrophil concentrations, will be needed to evaluate the extent to which these predictions hold across all five extant species in the Monotremata order. Additionally, we predicted the evolution of the lowest growth rate viruses in reservoir hosts of the orders Scandentia, Cetartiodactyla, Diprotodontia, and Dasyuromorphia, all of which demonstrated significantly low estimates for the magnitudes of constitutive immunity and two of which (Cetartiodactyla, Dasyuromorphia) also demonstrated significantly low estimates for tolerance of immunopathology.

### Estimating zoonotic virus virulence in spillover human hosts

After establishing optimal growth rates for viruses evolved in diverse reservoir host orders 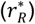, we subsequently modeled the corresponding ‘spillover virulence’ (*α*_*S*_) of these viruses following emergence into a human host. Zoonotic spillovers were modeled as acute infections in the human, and virulence was calculated while varying only the growth rate of the spillover virus 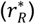 and the human tolerance of direct virus pathology (*T*_*vS*_) between viruses evolved in differing reservoir orders. We varied this last parameter, *T*_*vS*_, to account for any differences in virus adaptation to reservoir host immune systems that were not already captured in estimation of reservoir host *T*_*wR*_ and *g*_0*R*_. *T*_*vS*_ was thus computed as the inverse of the scaled time to MRCA for each mammalian reservoir host order from Primates (Fig 3E; S4 Fig, S1-S2 Tables), such that we estimated low human tolerance to viruses evolved in phylogenetically distant orders (e.g. monotreme and marsupial orders), and high human tolerance to viruses evolved in Primate and Primate-adjacent orders. In general, the modulating effects of *T*_*vS*_ did little to alter virulence rankings of zoonotic viruses from those predicted by raw growth rate 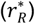 alone—with the highest spillover virulence predicted from viruses evolved in orders Monotremata and Chiroptera and the lowest spillover virulence predicted from viruses evolved in orders Cetartiodactyla and Scandentia (Fig 3D,G). Notably, modulating *T*_*vS*_ enhanced predictions of spillover virulence for some marsupial clades (Peramelemorphia, Dasyuromorphia, Diprotodontia) relative to eutherian orders with similar predicted 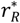 values. This resulted in dampened predicted spillover virulence for the eutherian order Perissodactyla as compared with marsupial orders Dasyuromorphia and Diprotodontia, despite lower predicted 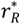 values for the latter two clades (Fig 3D,G; S5 Fig). Similarly, this elevated spillover virulence predictions for Peramelemorphia above Eulipotyphla, Afrosoricida, and Hyrocoidea, despite higher predicted 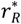 in the three eutherian orders (Fig 3D,G; S5 Fig).

### Comparing predictions of spillover virulence by reservoir order with estimates from the literature

Since parameter magnitudes estimated from life history traits were largely relative in nature, we scaled predictions of spillover virulence (*α*_*S*_) by reservoir order in relative terms to compare with estimates gleaned from the zoonosis literature (8) (Fig 3G, S5-S9 Figs). For eight reservoir orders (Chiroptera, Eulipotyphla, Primates, Carnivora, Rodentia, Diprotodontia, Perissodactyla, Cetartiodactyla), we were able to fit a simple linear regression comparing the relative virulence of zoonoses derived from each order as reported from case fatality rates in the literature (4,8) vs. those predicted by our nested modeling approach (S6,S8 Figs; S3 Table). Our nested modeling framework recovered available case fatality rate data well, yielding an R^2^ value of 0.57 in the corresponding regression of observed vs. predicted values (S6 Fig; S3 Table). Critically, we successfully recovered the key result from the zoonosis literature: bat-derived zoonoses yield higher rates of virus-induced mortality upon spillover to humans (*α*_*S*_) than do viruses derived from all other eutherian mammals (Fig 3G; S5-S8 Fig). In general, high estimates for *T*_*wR*_, *μ*_*R*_, and *g*_0*R*_, and low estimates for *T*_*vS*_ predicted high spillover virulence to humans (*α*_*S*_). Bats demonstrate uniquely long lifespans for their body sizes and uniquely enhanced constitutive immune responses (21) as compared with other taxa; when combined, as in our analysis, to represent high *T*_*wR*_ and high *g*_0*R*_, these reservoir host traits elevated predicted 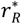 and *α*_*S*_ beyond all other eutherian orders (Fig 3D,G).

Evaluated against the data (4,8), our model overpredicted cross-order comparative virulence rankings for viruses evolved in order Eulipotyphla (Fig 3G; S6 Fig), largely as a result of our parameterization of high values for annual mortality rate (*μ*_*R*_) and correspondingly low values for tolerance of immunopathology (*T*_*wR*_) in this order. In addition, our model underpredicted virulence for Carnivora-derived viruses, based on within-host parameter estimates largely inverse to those recovered for Eulipotyphla (e.g. low *μ*_*R*_ and high *T*_*wR*_). We were able to resolve this underprediction of Carnivora virulence when excluding rabies lyssavirus from data comparisons (4,8) (S8 Fig); though most rabies zoonoses are sourced from domestic dogs, lyssaviruses are Chiropteran by origin (63), and viral traits responsible for rabies’ virulence may reflect its bat evolutionary history more than that of any intermediate carnivore host. Nonetheless, while excluding rabies from comparison improved recovery of literature-estimated relative virulence of Carnivora-derived viruses, it somewhat destabilized predictions for other orders, such that the overall performance of the model was largely equivalent as to when comparing against all available data (S8 Fig; R^2^ = 0.57).

In all cases, our model successfully reproduced estimates of significantly lower virulence incurred by zoonotic viruses evolved in Cetartiodactyla hosts than in all other orders considered for comparison (8). While previous work suggested that low observed virulence for Cetartiodactylan zoonoses might result from overreporting of avirulent zoonoses in domestic livestock with frequent human contact (8), our analysis indicates that viral zoonoses emerging from Cetartiodactylan hosts may truly be less virulent to humans. Our mechanistic framework demonstrates that reduced tolerance of immunopathology (manifest as shorter lifespans than predicted by body size) and limited constitutive immune responses identified in Cetartiodactylan hosts may drive the evolution of low growth rate viruses that cause correspondingly benign infections following zoonotic emergence. Further research into the extent to which Cetartiodactylans are impacted by immunopathology, as well as the degree to which low reported baseline neutrophil concentrations accurately reflect their innate immunity, is needed to evaluate these predictions. Quantification of the natural growth rates of Cetartiodactylan-evolved viruses could offer one means of testing this modeling framework.

Sensitivity analyses demonstrated that, as previously observed in Fig 2, parameters had unequal impacts on the resulting prediction of spillover virulence in our nested modeling framework (S9 Fig; S3 Table). Indeed, changes in *g*_0*R*_—followed by changes in *T*_*vS*_, and then *T*_*wR*_—had the greatest influence on the resulting predictions for *α*_*S*_. Likelihood optimization of *g*_0*R*_ while holding *μ*_*R*_, *T*_*wR*_, and *T*_*vS*_at order-specific values estimated from regression analyses of publicly available life history data greatly improved our recovery of spillover virulence as reported in the literature—yielding an R^2^ value of 0.99 from the corresponding regression of observed vs. predicted values (S9 Fig; S4 Table).

## Discussion

Our work provides a mechanistic insight into the biological processes that underpin cross-species patterns in the evolution of virus virulence—a major advance for efforts to evaluate zoonotic risk. Using a simple model of evolving virus in a reservoir host immune system, we successfully recapture patterns previously reported in the literature that document both the extreme virulence of viral zoonoses derived from bats and the surprising avirulence of viruses derived from Cetartiodactylan hosts (8). Notably, our nested modeling approach produces rank-ordered predictions of spillover virulence recovered in the literature (8), which are distinct from those that would result from a scaled inversion of phylogenetic distance alone. Additionally, our mechanistic approach allows us to make powerful predictions about the virulence of potential future viral spillovers, using general life history traits across mammals, including from orders for which zoonoses have not yet been reported (8). Our model indicates that we should anticipate the evolution of high growth rate viruses likely to cause virulence upon cross-species emergence from mammalian hosts with protracted lifespans for their body size (which we link to molecular mechanisms of immunopathological tolerance (49)), as well as from hosts with robust constitutive immune responses (36). While both immunopathological tolerance and constitutive immunity should drive the evolution of high growth rate viruses (Fig 2), only tolerance will do so in the absence of reservoir host pathology, thus highlighting its importance in driving observed variation in the virulence of viral zoonoses. Notably, while tolerance reduces pathogen-induced mortality for a single host, tolerant host populations (with limited checks on virus transmission) may demonstrate high pathogen prevalence. If imperfectly tolerant hosts still experience some virus-induced pathology, high prevalence can consequently elevate total population-level mortality for infected hosts—a phenomenon known as the ‘tragedy of tolerance’ (40,41). Reports of virus-induced mortality in bats are rare (64), suggesting that bat virus tolerance is likely very effective.

Intriguingly, our independent evaluations of mammalian life history traits associated with enhanced longevity (which we model as a proxy for immunopathological tolerance) and robust constitutive immunity highlight several mammalian orders—chiefly Chiroptera and Monotremata—which show synergy in both features, offering support for ongoing efforts to elucidate links between anti-aging molecular pathways and antiviral immunity (59,64). Further bolstering these hypotheses, we demonstrate the inverse synergy in order Cetartiodactyla, which exhibits shorter than predicted lifespans per body size but significantly reduced constitutive immune responses as compared with other mammals. Though exciting, our predictions should nonetheless be regarded with considerable caution, as taxon-specific insights resulting from our modeling framework are limited by a dearth of comparative data. For example, we characterize host tolerance of immunopathology based on deviations in lifespan as predicted for body size because more directly representative measures of, for example, mammalian antioxidant capacity, are not universally available. In addition, we model variation in reservoir host immunology almost exclusively based on variation in baseline neutrophil concentrations across mammalian orders, while holding constant many basic immunological parameters that almost certainly vary across taxa. Though crude, baseline neutrophil data nonetheless produce an estimate of enhanced constitutive immune responses for order Chiroptera, which is robustly supported by independent molecular work describing constitutive interferon expression in bat cells across multiple species (21,65,66).

All told, this analysis highlights a critical need for compilation of a more complete comparative immunological database that would enable quantification of the many additional within-host parameters (e.g. leukocyte activation rate, virus consumption rate, leukocyte mortality rate, and host tolerance of direct virus pathology) represented in our model. Our relative success in recapturing broad patterns in spillover virulence, despite data constraints, suggests that improvements in parameter estimation will almost certainly yield gains in our nuanced understanding of the process of cross-species virus emergence. For example, we model host tolerance of direct virus pathology as proportional to phylogenetic distance between reservoir and spillover host, but this term will likely also be modulated by virus tropism—presenting yet another mechanism by which viral adaptation to reservoir hosts could enhance spillover virulence. Indeed, coronavirus tropism is thought to be localized largely in the gastrointestinal tract for bat hosts (67,68): in our modeling framework, higher tolerance of direct virus pathology in this tissue should promote the evolution of higher growth rate viruses in bats, which are likely to cause virulence upon infection of more vulnerable tissues, such as the respiratory tract, in spillover hosts.

By summarizing within-host traits across mammalian orders, we generalize substantial within-clade diversity that likely also contributes to heterogeneous patterns in available data. Bats alone make up more than 1,400 species and account for some 20% of mammalian diversity (69). Only a subset of bats are long-lived (70), and the magnitude of constitutive immunity is known to vary across bat species (21,36,44), suggesting considerable variation in the parameters *μ*_*R*_, *T*_*wR*_, and *g*_0*R*_, which we here model universally across the entire Chiropteran order. Thus, we can expect considerable variation in the evolution of virulence for viruses evolved in diverse species within a given order—which we largely disregard here. As more data becomes available, our modeling approach could be fine-tuned to make more specific, species-level predictions of highly virulent disease risk.

Our analysis does not consider the probability of cross species viral emergence, or the potential for onward transmission of a spilled-over virus in a human host—both of which have been shown to correlate inversely with phylogenetic distance across mammals (4,8,9). Indeed, in keeping with trade-off theory, onward transmission of viruses following spillover is more commonly associated with low virulence infections (6,8,71,72), suggesting that reservoir orders highlighted here as potential sources for high virulence pathogens are not necessarily the same orders likely to source future pandemics. Nonetheless, the possibility for virus adaptation to gain transmission advantages in human hosts following spillover—as witnessed for Ebola (73) and SARS-CoV-2 (74)—should not be ignored.

Currently, our work emphasizes the uniqueness of bats as flying mammals and, in consequence, as long-lived, tolerant reservoirs for highly virulent viral zoonoses. For the first time, we offer a mechanism for the evolution of bat-derived viruses that demonstrate significant pathology upon spillover to non-bat, particularly human, hosts. In providing a theoretical framework to explain this phenomenon, we generate a series of testable questions and hypotheses for future comparative immunological studies, to be carried out at *in vitro* and *in vivo* scales. Empirical work should aim to measure rates of immune cell activation, growth, and mortality across diverse mammalian orders and determine whether natural virus growth rates are truly higher when evolved in bat hosts. Additional studies should test whether anti-inflammatory mechanisms in bat cells of different tissues are equally effective at mitigating virus-induced pathology and immunopathology and whether comparative taxonomic predictions of virus tolerance, resistance, and virulence evolution apply to non-viral pathogens, too. In light of the emergence of SARS-CoV-2, the field of bat molecular biology has echoed the call for mechanistic understanding of bat immunology (75), and the NIH has responded (76), soliciting research on the development of tools needed to test the predictions outlined here. We offer a bottom-up mechanistic framework enabling the prediction of emerging virus virulence from the basic immunological and life-history traits of zoonotic hosts.

## Methods

### Evolutionary framework

#### Population-level dynamics

To evaluate the selective pressures that drive the evolution of cross-species virus virulence, we first sought to derive an equation for an optimal virus growth rate expressed in within-host parameters specific to the life history of the reservoir host. To this end, and in keeping with classic examples of viral maintenance in reservoir populations (77–83), we first modeled the dynamics of a persistent infection (*I*_1_) in a hypothetical reservoir host, allowing for the introduction of a rare mutant virus strain which generates distinct infections in that same host (*I*_2_) (see S1 File for more detailed methodology and derivations):

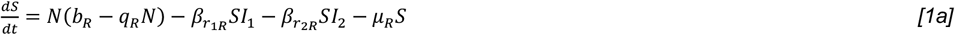

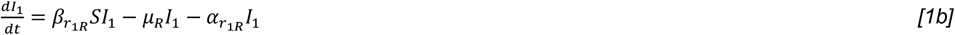

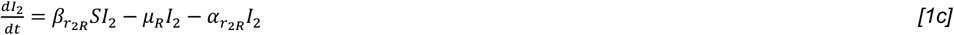

Here, *N* corresponds to the total host population and *S* to all hosts susceptible to infection, such that *N* = *S* + *I*_1_ + *I*_2_. We assumed that hosts are born at rate *b*_*R*_ and die of natural death at rate *μ*_*R*_, where *b*_*R*_ > *μ*_*R*_. We further assumed that all hosts are born susceptible and that population density is regulated via a crowding term (*q*_*R*_) applied to the birth rate. The subscript “_R_” on the birth, death, and crowding terms emphasizes that these rates are specific to the reservoir host. Because we aimed to model the evolution of rates that link to within-host dynamics for the reservoir host, we further represented transmission and virulence as functions of the virus causing the infection (respectively, 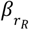 and 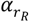) where *r*_*R*_ denotes the intrinsic virus growth rate and is represented distinctly for both the endemic (*r*_1*R*_) and mutant (*r*_2*R*_) virus strains.

With the use of model [1], we performed an evolutionary invasion analysis (see S1 File) and concluded that the virus should evolve to maximize the ratio of transmission over infection duration. We refer to this ratio as the invasion fitness:

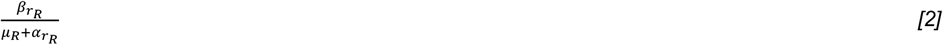

#### Within-host dynamics

Next, to evaluate the within-host selective conditions underpinning the evolution of the virus growth rate (*r*_*R*_), we sought to express the between-host parameters of transmission 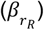 and virulence 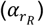 in within-host terms, using a nested modeling approach. To this end, we established a simple within-host model representing the dynamics of infection *within* each *I*_1_and *I*_2_ host as outlined above. We followed Alizon and Baalen 2005 (45) to adapt a class of Lotka-Volterra predator-prey-like within-host models (reviewed in (84)), to overcome some constraints of basic predator-prey models of the immune system, chiefly *(i)* by allowing leukocytes to circulate in the absence of infection and *(ii)* by scaling leukocyte growth with virion density, independent of direct leukocyte-virion contact (45,85,86). This resulted in the following model, which demonstrates interactions between the virus population (V_R_) and the leukocyte population (L_R_) within each infected reservoir host:

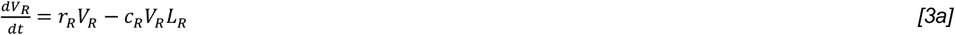

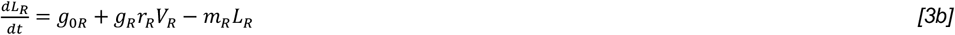

Here, *r*_*R*_ corresponds to the intrinsic virus growth rate,*c*_*R*_ corresponds to the attack efficacy of the immune system upon contact with the virus, and *g*_*R*_ signifies the recruitment rate of immune cells scaled to the virus growth rate. The parameter, *g*_0*R*_, describes the constitutive, baseline leukocyte recruitment in the absence of infection, and *m*_*R*_ gives the natural leukocyte death rate, all in the reservoir host.

Building from above, we expressed rates of population-level transmission and virulence known to depend on within-host dynamics (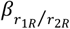 and 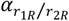) in terms of their within-host components, assuming that within-host dynamics are fast relative to host population-level dynamics and, therefore, converge to the endemic equilibrium (i.e. *m*_*R*_*r*_*R*_ > *c*_*R*_*g*_0*R*_).

In line with previous work, we assumed transmission to be a linear function of viral load (45), which we represented as:

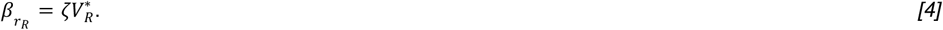

where *ζ* corresponds to a scaling term equating viral load to transmission. We additionally assumed that:

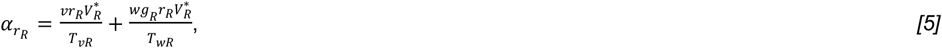

by which infection-induced host mortality (‘virulence’) is modeled to result from both virus-induced pathology (a function of the intrinsic virulence of the parasite, *v*, multiplied by the parasite growth rate 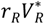) and immunopathology—which we modeled as proportional to the leukocyte growth rate (*g*_*R*_) multiplied by the virus growth rate 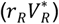 multiplied by the parasite’s intrinsic propensity for host immune antagonism (*w*). The terms *T*_*vR*_ and *T*_*wR*_ correspond to host tolerance of virus-induced pathology and immunopathology, respectively, under assumptions of “constant” tolerance, by which both virus pathology and immunopathology are reduced by a constant proportion across the course of infection. For constant tolerance, we assumed that *T*_*vR*_ > 1 and *T*_*wR*_ > 1. See S1 File for detailed derivations under assumptions of “complete” tolerance, whereby virus pathology and immunopathology are completely eliminated up to a threshold value, beyond which pathology scales proportionally with virus and immune cell growth.

Now, we rewrote the equations for transmission (*[4]*) and virulence (*[5]*) in purely within-host terms:

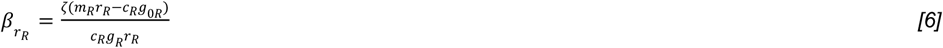

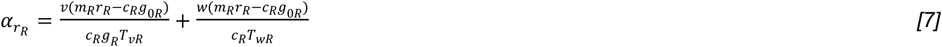

Using these within-host expressions, we then recomputed the expression for invasion fitness (*[2]*) in within host terms and determined the optimal intrinsic virus growth rate, which is an evolutionarily stable strategy (ESS) (see S1 File):

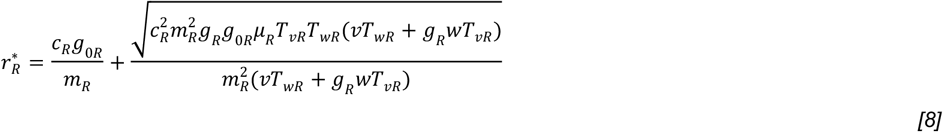

In Fig 2, we explore the sensitivity of 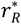, and the corresponding reservoir host population-level transmission 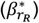 and virulence 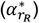 at that growth rate, across varied values for its within-host component parameters.

#### Cross-species dynamics

We next sought to derive an expression to explore the consequences of spillover of a virus evolved to its optimal growth rate in a reservoir host population – which we termed 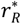 – following cross-species emergence into the human population. Contrary to its established persistent infection in the reservoir host, we assumed that such a virus would produce an acute infection in the spillover host. To model the dynamics of this acute spillover, we borrowed from Gilchrist and Sasaki 2002 (47), who developed a within-host parasite-leukocyte model in which an immortal leukocyte successfully eradicates the parasite population to near-zero. We modified their acute model to be more comparable to our chronic infection model (*[3a]/[3b])* and to reflect our differing notation for within-host dynamics.

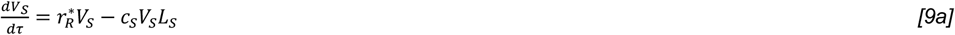

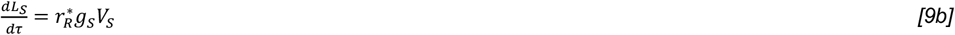

Here, 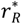 represents the reservoir-evolved virus growth rate, but all other terms reflect within-host conditions of the spillover host: *V*_*S*_ and *L*_*S*_ correspond, respectively, to the spillover host virus and leukocyte populations, *c*_*S*_ is the virus consumption rate upon contact with leukocytes in the spillover host, and *g*_*S*_ is the growth rate of the spillover host leukocyte population in response to virus. We expressed the model in units of *τ*, which we assumed to be short in comparison to the *t* time units of dynamics in the reservoir host population. Rather than solving for virus and leukocyte populations at equilibrium (as in the reservoir host population), we followed Gilchrist and Sasaki 2002 (47) to instead derive an expression for the virus population at the peak of infection 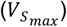, where we anticipated maximum pathology for the spillover host. We then extended this prior work to generate an expression for the average viral load 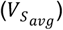 in the spillover host across the timecourse of acute infection, which we can use in comparisons of spillover host virulence with reported case fatality rates for zoonoses in the literature.

If we divide equation *[9a]* by equation *[9b]* we derive a simple time-independent relationship between virus and leukocyte density:

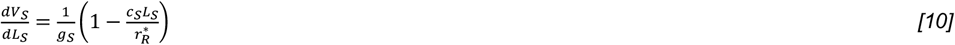

If we then let *L*_*S*_(0) = 1, *V*_*S*_(0) = 1, assuming that both virus and leukocyte populations will be small at *τ* = 0, we can integrate *[10]* to establish the following relationship between virus and leukocyte:

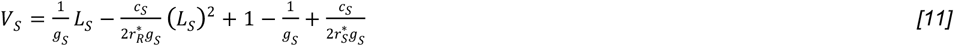

Notice that the above *[11]* is simply a quadratic equation (see *[12]* below):

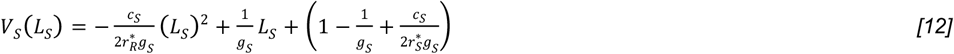

Now, we can take the derivative of equation *[12]* to formulate an expression for 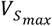, the maximum viral load, which should precede the end of acute infection and the point of host recovery at the maximum duration of infection:

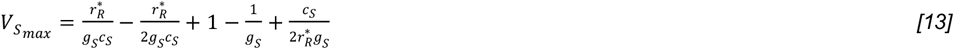

Then, extending previous work (47), we calculated the average viral load by taking the integral of equation *[12]* from *V*_*S*_(0)to 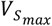 and dividing by the duration of that interval. From this exercise, we expressed the average value of equation *[12]* as:

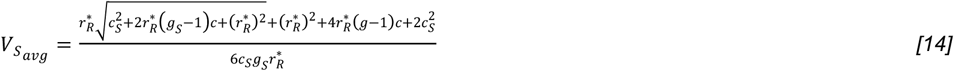

Now, with this established expression for 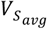, we adapted equation *[7]* to reflect the acute within-host dynamics of “spillover virulence”, which we modeled (as before) as a combination of mortality induced from direct virus pathology and from immunopathology. As in the reservoir host system, we also modeled the mitigating impact of tolerance on the two mechanisms of virulence:

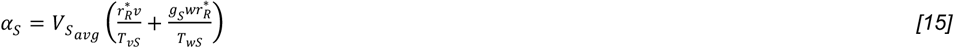

This yielded the above expression for ‘spillover virulence’. Here, the growth rate of the virus is expressed at its evolutionary optimum evolved in the reservoir host 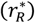, and the virus’s intrinsic virulence (*v*) and propensity to elicit an inflammatory immune response (*w*) remained unchanged from one host to the next. Inspired by findings in the literature that report higher virulence in cross-species infections between hosts separated by larger phylogenetic distances (8–12), we modeled spillover host tolerance of virus-induced pathology (*T*_*vS*_) as a decreasing function of increasing phylogenetic distance between the reservoir and secondary host. All other immune-related parameters assumed characteristics of the spillover host: *g*_*S*_ is the spillover host’s leukocyte growth rate, and *T*_*wS*_ corresponds to the spillover host’s tolerance of immunopathology.

### Order-specific estimates for optimal virus growth rates evolved in reservoir hosts

Next, we sought to develop estimates for the optimal virus growth rate 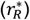 expected to evolve across a suite of diverse mammalian reservoir orders. Because within-host immunological data needed to quantify within-host parameters underpinning the expression for 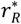 are largely lacking, we chose to proxy a few key within-host parameters from well-described allometric relationships for mammalian life history data. As such, we here focused on three key parameters for which data could be gleaned from the literature: the reservoir host mortality rate (*μ*_*R*_), the reservoir host tolerance of immunopathology (*T*_*wR*_), and the magnitude of reservoir host constitutive immunity (*g*_0*R*_).

To generate order-level summary terms for *μ*_*R*_, we fit a simple linear regression to the response variable of the inverse of maximum lifespan (in days) with a single categorical predictor of host order, using data from Jones et al. 2009 (54) and Healy et al. 2014 (55) that spanned 26 mammalian orders and 1060 individual species (Fig 3A). This first model thus took the form:

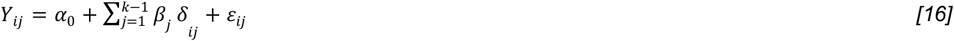

where *Y*_*ij*_ corresponds to natural mortality rate observations for *i* species belonging to *j* orders; *α*_0_ is the overall intercept; *k* indicates the total number of available orders (here, 26); *β*_*j*_ is an order-specific slope; *δ*_*ij*_ indicates observations of *i* species grouped into *j* orders; and *ε*_*ij*_ is a normally distributed error term corresponding to species *i* within order *j*. We estimated *μ*_*R*_by simply generating predictions from this fitted model across all 26 mammalian orders represented in our dataset (Fig 3C; S2 Table).

We next used the same dataset of 1060 species grouped into 26 mammalian orders to generate order-level estimates for the reservoir host tolerance of immunopathology (*T*_*wR*_). Here, we fit a linear mixed effects regression to the log_10_ relationship of lifespan (in years) as predicted by body mass (in grams). This second model took the form:

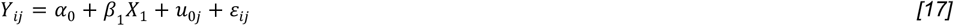

where *Y*_*ij*_ corresponds to the log_10_ value of record of maximum lifespan (in years) for *i*species belonging to *j* orders; *α*_0_ is the overall intercept; *β*_1_ indicates the slope of the fixed predictor of log_10_ mass (in grams), here represented as *X*_1_; *u*_0*j*_ is the order-specific random intercept; and *ε*_*ij*_ is a normally distributed error term corresponding to species *i* within order *j*. To estimate *T*_*wR*_, we extracted the relative partial effects of mammalian order on maximum lifespan per body size, then rescaled these effects between 1 and 2 for assumptions of constant tolerance and between 0 and 1 for assumptions of complete tolerance (S1-S2 Tables). We justified this approach based on literature that highlights links between anti-aging molecular pathways that promote longevity and those that mitigate immunopathology (11,44,57–61).

Finally, to generate order-level summary terms for the magnitude of constitutive immunity (*g*_0_), we fit another linear mixed effects regression to the relationship between the predictor variables of log_10_ mass (in g) and basal metabolic rate (BMR, in W) and the response variable of log_10_ baseline neutrophil concentration (in 10^9^ cells/L), which offers an approximation of a mammal’s constitutive innate immune response. BMR data for this model were derived from Jones et al. 2009 (54) and Healy et al. 2014 (55), while neutrophil concentrations were obtained from zoo animal data presented in the Species360 database (53); prior work using this database has demonstrated scaling relationships between body size and immune cells across mammals (52). Paired neutrophil and BMR data were limited to just 19 mammalian orders and 144 species. Our model also included a random effect of host order, resulting in the following form:

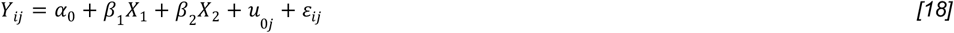

where *Y*_*ij*_ corresponds to the log_10_ baseline neutrophil concentrations (in 10^9^ cells/L) for *i* species belonging to *j* orders; *α*_0_ is the overall intercept; *β*_1_ indicates the slope of the fixed predictor of log_10_ mass (in grams), here represented as *X*_1_; *β*_2_ indicates the slope of the fixed predictor of BMR (in W), here represented as *X*_2_; *u*_0*j*_ is the order-specific random intercept; and *ε*_*ij*_ is a normally distributed error term corresponding to species *i* within order *j*. Using a similar approach as that employed above for *T*_*wR*_, we estimated *g*_0_ by extracting the relative partial effects of mammalian order on neutrophil concentration per mass-specific BMR, then rescaled these effects between 0 and 1. Because we estimated significantly positive partial effects between the orders Chiroptera and Monotremata and baseline neutrophil concentration (Fig 3C), this generated correspondingly high estimates of *g*_0*R*_ for these two orders (S4 Fig, S1 Table).

Using these order-level summary terms for *μ*_*R*_, *T*_*wR*_, and *g*_0*R*_, we then generated an order-level prediction for 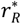 across all mammalian host orders, following equation *[8]* (Fig 3D). All other parameters (*c*_*R*_, *g*_*R*_, *m*_*R*_, *w, v*, and *T*_*vR*_) were held constant across all taxa at values listed in Table 1.

### Estimating zoonotic virus virulence in spillover human hosts

Once we had constructed an order-level prediction for 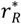, we finally sought to explore the effects of these reservoir-evolved viruses upon spillover to humans, following equation *[15]*. As when computing virulence for the reservoir host population, we held immunological parameters,*c*_*S*_, *g*_*S*_, and *m*_*S*_ constant in humans (at the same values listed above) due to a lack of informative data to the contrary. Then, to generate an order-level estimate for virus virulence incurred on humans (*α*_*S*_), we combined our order-level predictions for 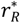 with order-specific values for *T*_*vS*_, the human tolerance of an animal-derived virus, which we scaled such that viruses derived from more closely-related orders to Primates were more easily tolerated in humans. Specifically, we represented *T*_*vS*_ as the scaled inverse of the cophenetic phylogenetic distance of each mammalian order from Primates. No summarizing was needed for *T*_*vS*_ because, following Mollentze and Streicker 2020 (10), we used a composite time-scaled reservoir phylogeny derived from the TimeTree database (56), which produced a single mean divergence date for all clades. Thus, all species within a given mammalian order were assigned identical times to most recent common ancestor with the order Primates. To convert cophenetic phylogenetic distance into reasonable values for *T*_*vS*_, we divided all order-level values for this distance by the largest observed (to generate a fraction), then subtracted that fraction from 2 for assumptions of constant tolerance (yielding *T*_*vS*_ estimates ranging from 1-2) and from 1 for assumptions of complete tolerance (yielding *T*_*vS*_ estimates ranging from 0-1).

We then combined reservoir host order-level predictions for 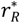 with these estimates for human tolerance of virus pathology (*T*_*vS*_) to generate predictions of the expected spillover virulence of viral zoonoses emerging from the 19 mammalian orders for which we possessed complete data across the four variable parameters (*μ*_*R*_, *T*_*wR*_, *g*_0*R*_, and *T*_*vS*_). We scaled these *α*_*S*_ estimates—and their 95% predicted confidence intervals—in relative terms (from 0 to 1) to compare with estimates from the literature.

### Comparing predictions of spillover virulence by reservoir order with estimates from the literature

To compare estimates of spillover virulence generated from our nested modeling approach (*α*_*S*_) with estimates from the literature, we followed Day 2002 (87) to convert case fatality rates of viral zoonoses in spillover human hosts (*CFR*_*S*_) reported in the literature (8) to empirical estimates of *α*_*S*_ for each mammalian order, using data on the duration of human infection (*D*_*IS*_) for each viral zoonosis. For this purpose, we collected data on *D*_*IS*_ by searching the primary literature; raw data for infection durations and associated references are reported in our publicly available GitHub repository (github.com/brooklabteam/spillover-virulence). Briefly, Day 2002 (13) notes that the equation for case fatality rate in the spillover host (*CFR*_*S*_) takes the form:

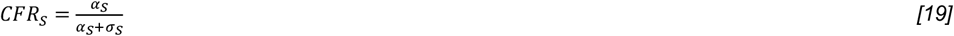

where *σ*_*S*_ corresponds to the recovery rate from the spilled-over-virus in the human host. We also note that the total duration of infection, *D*_*IS*_, is given by:

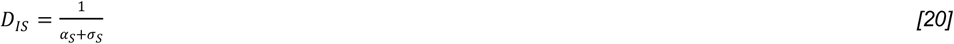

Therefore, we translated human case fatality rates of viral zoonoses (*CFR*_*S*_) into estimates of spillover virulence (*α*_*S*_) using the following equation:

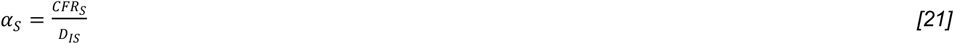

To obtain a composite order-level prediction for *α*_*S*_ from *D*_*IS*_ and *CFR*_*S*_ as reported in (8), we adopted the best-fit generalized additive model (GAM) (88) used by the authors in (8) to summarize *CFR*_*S*_ by order and, here, applied it to *α*_*S*_ estimates converted from *CFR*_*S*_ reported in the literature (S2 Table). We then summarized *α*_*S*_ at the order-level from the fitted GAM, excluding the effects of viral family. This yielded a literature-derived value for *α*_*S*_ across disparate mammalian orders, against which to compare predictions from our nested modeling approach. Because the majority of the within-host parameters underpinning *α*_*S*_ predictions in our nested modeling approach (*μ*_*R*_, *T*_*wR*_, *g*_0*R*_, and *T*_*vS*_) were quantified only in a relative fashion, we were unable to compare direct magnitudes of virulence (e.g. in terms of host death per unit time). To account for this, we rescaled *α*_*S*_ estimates from both our nested modeling approach and from the empirical literature from 0 to 1 and compared the relative rank of virulence by mammalian order instead. We predicted *α*_*S*_ for 19 discrete mammalian orders, though data from the literature were available for only 8 orders against which to compare.

Finally, because bats are the most likely ancestral hosts of all lyssaviruses (63), and it may be more appropriate to class carnivores as ‘bridge hosts’ for rabies, rather than reservoirs (as the authors discuss in Guth et al. 2022 (8)), we recomputed literature-derived estimates of relative *α*_*S*_ by fitting the same GAMs (S2 Table) to a version of the dataset excluding all entries for rabies lyssavirus. We rescaled these new estimates of spillover virulence between 0 and 1 and compared them again to predictions from our nested modeling approach.

To quantitatively evaluate the extent to which our nested model accurately recaptured estimates of relative spillover virulence (*α*_*S*_) recovered from the literature, we fit a simple linear regression to the relationship between observed and predicted virulence across the eight mammalian orders for which we possessed comparative data (S6, S8 Figs; S3 Table). We then conducted a sensitivity analysis to determine the extent to which our predictions of *α*_*S*_ across reservoir orders could be modulated by changes in the parameters we estimated from the literature. Because reservoir host natural mortality rate (*μ*_*R*_) had minimal influence on optimal virus growth rates 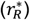 – and because we assumed mortality data to be the most likely to be accurately reported in the literature – we focused our analysis on the extent to which variation in estimation of reservoir host tolerance of immunopathology (*T*_*wR*_), reservoir host magnitude of constitutive immunity (*g*_0*R*_), and spillover host tolerance of reservoir-derived virus pathology (*T*_*vS*_) impacted the resulting prediction for spillover virulence (*α*_*S*_) (S9 Fig; S3 Table). To this end, we quantified changes in the maximum likelihood fit of our nested model’s predicted spillover virulence to the literature-derived data while profiling each of these three parameters (*T*_*wR*_, *g*_0*R*_, *T*_*vS*_) in turn across a range of reasonable values—both while holding all other parameters at constant values across all taxa and at heterogenous values estimated from regression analysis presented here in the main text (S9 Fig; S3 Table). To quantify the impact of this profiling on the overall recovery of relative spillover across our dataset, we refit linear regressions of observed vs. predicted spillover virulence to each new data subset generated from individual profiling of variable parameters (S9 Fig; S3 Table).

## Acknowledgments

The authors thank the Boots Lab at UC Berkeley for helpful comments on this manuscript.

## Supporting Information

**S1 Fig. Pairwise invasibility plots demonstrate optimal viral growth rate, under low and high tolerance conditions**. Invading growth rates (*r*_*R*2_) will displace resident growth rates (*r*_*R*1_) at values indicated by the shaded regions. Reservoir host tolerance of immunopathology (*T*_*wR*_) and tolerance of direct virus pathology (*T*_*vR*_) are both modeled as low in left column (0.5 and 10 for row 1 and 2, respectively) and high in right column (0.97 and 100 for row 1 and 2), assuming a complete (row 1) or a constant form (row 2). For this visualization, *r*_*R*1_and *r*_*R*2_ span from 3.18 to 3.5. All other parameters involved in computation of 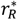 (see S1 File equations *[25]* and *[26]*) were fixed at values listed in Table 1 (main text).

**S2 Fig. Optimal virus growth rates—and subsequent spillover virulence—vary across reservoir host immunological and life history parameters**. Figure replicates Fig 2 (main text) under assumptions of complete tolerance. Rows (top-down) indicate the evolutionarily optimal within-host virus growth rate 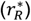 and the corresponding transmission rate 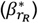, and virus-induced mortality rate 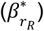 for a reservoir host infected with a virus at 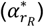. The bottom row then demonstrates the resulting virulence (*α*_*S*_) of a reservoir-optimized virus evolved to 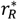 upon nascent spillover to a novel, secondary host. Columns demonstrate the dependency of these outcomes on variable reservoir host parameters: background mortality rate (*μ*_*R*_), extent of constitutive immunity (*g*_0*R*_), leukocyte activation rate upon viral contact (*g*_*R*_), virus consumption rate by leukocytes (*c*_*R*_), leukocyte mortality rate (*m*_*R*_). Darker colored lines depict outcomes at higher values for reservoir host tolerance of virus pathology (*T*_*vR*_,red) or immunopathology (*T*_*wR*_,blue), assuming no tolerance of the opposing type. Heat maps demonstrate how *T*_*vR*_ and *T*_*wR*_interact to produce each outcome. Outcome ranges differ between lineplots (y-axes) and heatmaps (scalebars). Parameter values are listed in Table 1 (main text).

**S3 Fig. Spillover host tolerance of direct virus pathology and immunopathology modulate resulting virulence of spilled-over viruses**. Virulence of spilled-over virus (*α*_*S*_) across a range of values for two mechanisms of tolerance in the spillover host: tolerance of direct virus pathology (*T*_*vS*_, left column) and tolerance of immunopathology (*T*_*wS*_, right column). Results are expressed under assumptions of constant tolerance (top panels) and complete tolerance (bottom panels). In main text results, *T*_*wS*_ is held constant for all predictions of spillover virulence but *T*_*vS*_ is varied proportionally to the inverse time to MRCA between reservoir and spillover host.

**S4 Fig. Parameter estimates for life history traits across mammalian orders**. Model parameter estimates for (A) reservoir-host background mortality (*μ*_*R*_), (B) tolerance of immunopathology (*T*_*wR*_) (left y-axis: constant tolerance assumptions; right y-axis: complete tolerance assumptions), (C) magnitude of constitutive immunity (*g*_0*R*_), and (D) magnitude of human tolerance of virus pathology for a virus evolved in a disparate mammalian reservoir (*T*_*vS*_). Estimates are derived from (A) linear model predictions of maximum lifespan at the order level, (B) the scaled effect of order on a linear mixed model prediction of lifespan per body size, (C) the scaled effect of order on a linear mixed model prediction of neutrophil concentration for mass-specific BMR, and (D) the magnitude of human tolerance of virus pathology for a virus evolved in a disparate mammalian reservoir (*T*_*vS*_), corresponding to data presented in Fig 3E (main text). Default parameter values involved in the estimation process are summarized in Table 1 (main text), and estimated parameters and corresponding 95% confidence intervals by standard error are presented in S1 Table. See main text Methods and our open-source GitHub repository for a detailed walk-through of the parameter estimation process.

**S5 Fig. Optimal virus growth rates** 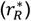 **and subsequent spillover virulence (***α*_*S*_**) for viruses evolved in diverse mammalian reservoirs**. Figure replicates Fig 3D and Fig 3G from the main text, here under assumptions of complete tolerance. Panel (A) depicts optimal 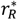 across 19 mammalian orders, for which we were able to estimate order-level specific values for the three within-host reservoir parameters which we varied in our analysis (*μ*_*R*_, *T*_*wR*_, and *g*_0*R*_; visualized in S4 Fig), while panel (B) depicts the resulting estimation of relative spillover virulence (*α*_*S*_), which also relies on order-specific values for the spillover host tolerance of direct virus pathology (*T*_*vS*_). Taxa in panels (A) and (B) are arranged in descending order from highest to lowest predicted values for, respectively 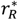 and *α*_*S*_. This order varies slightly from panel (A) to (B), as highlighted by alluvial flows and discussed in the main text. See S1 Table for order-level values for 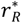, *μ*_*R*_, *T*_*wR*_, *g*_0*R*_, and *T*_*vS*_ and Table 1 (main text) for all other parameters involved in calculation of 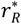 and *α*_*S*_.

**S6 Fig. Comparison of human spillover virulence (***α*_*S*_ **) as observed in the literature vs. predicted from nested modeling framework**. Figure plots observed vs. predicted spillover virulence for eight orders from Fig 3G (main text) for which case fatality rates from corresponding zoonoses are reported in the literature (8). Panel (A) compares nested modeling predictions under assumptions of constant tolerance with those from the literature, while panel (B) does the same under assumptions for complete tolerance. In both cases, a fitted linear regression and corresponding R^2^ value is shown as a quantitative evaluation of model fit to the data. Dashed lines give the residual of each data point from the regression line.

**S7 Fig. Comparison of relative human spillover virulence (***α*_*S*_**) predictions for zoonoses from nested life history model with estimates from the literature, excluding rabies**. Figure replicates Fig 3G (main text), respectively, under assumptions of (A) constant and (B) complete tolerance but excluding rabies lyssavirus from the zoonotic data (right-half of panels). Rank-order predictions of virulence are more consistent with order Carnivora further down in the rankings. As in Fig 3G, order-specific parameter values for 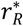, *μ*_*R*_, *T*_*wR*_, *g*_0*R*_, and *T*_*vS*_ are listed in S1 Table; all other parameters involved in calculation of *α*_*S*_ are listed in Table 1 (main text).

**S8 Fig. Comparison of human spillover virulence (***α*_*S*_ **) as observed in the literature (excluding rabies) vs. predicted from nested modeling framework**. Plot recapitulates S6 Fig exactly, but comparisons are drawn from case fatality rates reported in the literature but excluding rabies lyssavirus, which is often classed as a Carnivora-derived virus, though its evolutionary origins are found in bats. Removal of rabies improves estimates of virulence for Carnivora-derived zoonoses as compared with the complete dataset, but resulting linear regression offers no better fit to the entire dataset than previously shown in S6 Fig.

**S9 Fig. Sensitivity analysis of individual parameter influence on nested model fit to observed data**.Figure replicates S6 Fig in part with observed spillover virulence (*α*_*S*_) from case fatality rates reported in the literature (8) depicted on the x-axis and predictions from nested modeling framework on the y-axis. In all panels, circles correspond to nested modeling predictions of spillover virulence using parameter values recovered from regression analysis of publicly available life history data as presented in the main text, replicating points from S6 Fig. Projections from nested modeling approach assuming constant tolerance are shown in the top panels and complete tolerance in the bottom. In lieu of life history-derived parameter values, squares show *α*_*S*_ estimates from nested model using universal values across order for all parameters excepting the profiled optimum of the parameter corresponding to the column in question (*T*_*wR*_, *T*_*vS*_, or *g*_0*R*_). When not profiled, *T*_*wR*_=1.5 (constant) and 0.5 (complete); *T*_*vS*_=1.5 (constant) and 0.5 (complete); and *g*_0*R*_=0.5 for simulations resulting in square points. Finally, triangles give *α*_*S*_ estimates from nested model approach using parameters generated by profiling the optimum of the parameter corresponding to the column in question (*T*_*wR*_, *T*_*vS*_, or *g*_0*R*_) and using regression analysis values from the literature for other variable parameters (S1 Table). Lines and corresponding R^2^ values signify the fit of a simple linear regression of observed vs. predicted *α*_*S*_ across all mammalian orders, where predicted values are generated from nested modeling approach using: linear regression analysis of life history data for *T*_*wR*_, *T*_*vS*_, and *g*_0*R*_ (solid line, red); profiling *T*_*wR*_, *T*_*vS*_, or *g*_0*R*_while holding constant all other parameters across orders (thin dashed line); and profiling *T*_*wR*_, *T*_*vS*_, or *g*_0*R*_while using linear regression estimates from life history data for parameters not being profiled (thick dashed line).

**S1 Table. Order-specific parameter values for within-host nested model**.

**S2 Table. Model summary outputs for within-host parameter estimation**.

**S3 Table. Sensitivity analysis and comparison of influence of individual parameter estimates on overall fit of nested model to data**.

**S1 File. Supplementary Information Text**.

