## Supplementary Appendix for "Reservoir host immunology and life history shape virulence evolution in zoonotic viruses"

### Supplementary Information Text

#### S1 Text. Population-level dynamics and $R_0$

To begin, we follow Miller et al. (2006) (1), assuming a simple population dynamics system which allows for infection-induced mortality. However, since we are interested in examining virus evolution in reservoir hosts, we first assume that hosts maintain persistent lifelong infections following initial exposure.

We assume that hosts are born at rate $b_R$ and die of natural death at rate $\mu_R$, where $b_R > \mu_R$. All hosts are assumed to be born susceptible and population density is regulated via a crowding term ($q_R$). The subscript "R" on the birth, death, and crowding terms emphasizes that these rates are specific to the reservoir host. Because our goal in this endeavor is to allow for the evolution of rates that link to within-host dynamics (Text S3), we represent transmission and virulence as functions of the virus causing the infection respectively, ($\beta_{r_R}$ and $\alpha_{r_R}$) where $r_R$ denotes the intrinsic virus growth rate. Our system is described by the following system of ordinary differential equations (ODE), where 'S' represents hosts susceptible to infection and 'I' represents infected hosts and ($N = S + I$):

$$\frac{dS}{dt} = N(b_R - q_R N) - \beta_{r_R} SI - \mu_R S \qquad [1a]$$

$$\frac{dI}{dt} = \beta_{r_R} SI - \mu_R I - \alpha_{r_R} I \qquad [1b]$$

There exists both a disease-free equilibrium (DFE) at $(I^*_{DFE}, S^*_{DFE}) = (0, \frac{b_R - \mu_R}{q_R})$, as well as an endemic infection equilibrium (EE), $(I^*_{EE}, S^*_{EE})$ (see below, *[2a], [2b]*).

$$S^*_{EE} = \frac{\alpha_{r_R} + \mu_R}{\beta_{r_R}} \qquad [2a]$$

$$I^*_{EE} = \frac{\sqrt{\beta_{r_R}\left[\beta_{r_R}\left(b_R - \mu_R - \alpha_{r_R}\right)^2 + 4\,\alpha_{r_R} q_R\left(\mu_R + \alpha_{r_R}\right)\right]} + \beta_{r_R}\left(b_R - \mu_R - \alpha_{r_R}\right) - 2q_R\left(\alpha_{r_R} + \mu_R\right)}{2\,\beta_{r_R} q_R} \qquad [2b]$$

From *[1]*, we can then formulate the Jacobian matrix, $J$, to decipher the local stability of the DFE:

$$J = \begin{bmatrix} b_R - 2q_R S - 2q_R I - \beta_{r_R} I - \mu_R & b_R - 2q_R S - 2q_R I - \beta_{r_R} S \\ \beta_{r_R} I & \beta_{r_R} S - \mu_R - \alpha_{r_R} \end{bmatrix} \qquad [3a]$$

Next, we evaluate the Jacobian at the DFE:

$$J_{DFE} = \begin{bmatrix} -b_R + \mu_R & -b_R + 2\mu_R - \frac{\beta_{r_R}(b_R - \mu_R)}{q_R} \\ 0 & \frac{\beta_{r_R}(b_R - \mu_R)}{q_R} - \mu_R - \alpha_{r_R} \end{bmatrix} \qquad [3b]$$

The DFE is locally unstable when at least one of the eigenvalues of the Jacobian is non-negative. The eigenvalues can be read off the diagonal ($J_{DFE}$ is upper triangular).

Clearly, the first eigenvalue is negative since $b_R > \mu_R$: $\mu_R - b_R < 0$. A little algebra will establish that the second eigenvalue is negative when:

$$R_0 = \frac{\beta_{r_R}(b_R - \mu_R)}{q_R(\mu_R + \alpha_{r_R})} < 1 \qquad [4]$$

Therefore, if $R_0 > 1$, the DFE is no longer locally asymptotically stable, resulting in an increase in infected hosts within the population.

When $R_0 > 1$, the EE is globally asymptotically stable in the interior of $\Omega = \{(S, I) \in R_+^2 \mid S \geq 0, I \geq 0, S + I \leq \frac{b_R - \mu_R}{q_R}\}$. We demonstrate this by first establishing the local asymptotic stability of the EE, then by determining that there are no limit cycles in the region, $\Omega$.

If we evaluate the Jacobian, once again, at the EE (which we can refer to now as $J_{EE}$), we will find that the trace *(Trace($J_{EE}$))* of that matrix is negative:

$$Trace(J_{EE}) = \frac{-\left(2q_R + \beta_{r_R}\right)\sqrt{\beta_{r_R}\left[\beta_{r_R}\left(b_R - \mu_R - \alpha_{r_R}\right)^2 + 4\,\alpha_{r_R} q_R\left(\mu_R + \alpha_{r_R}\right)\right]} - \beta_{r_R}\left[\beta_{r_R}\left(b_R - \mu_R - \alpha_{r_R}\right) - 2q_R\left(\mu_R + \alpha_{r_R}\right)\right]}{2q_R\beta_{r_R}} < 0 \qquad [5]$$

Note that $b \geq \mu + \alpha_r$ (a necessary condition for the existence of the EE). In addition, the determinant of the $J_{EE}$ is positive:

$$Det(J_{EE}) = \frac{\sqrt{\beta_{r_R}\left[\beta_{r_R}\left(b_R - \mu_R - \alpha_{r_R}\right)^2 + 4\,\alpha_{r_R} q_R\left(\mu_R + \alpha_{r_R}\right)\right]}\left[1 - 2q_R\left(\mu_R + \alpha_{r_R}\right)\right] + \beta_{r_R}\left(b_R - \mu_R - \alpha_{r_R}\right)}{2q_R\beta_{r_R}} > 0 \qquad [6]$$

Note that $0 < q_R \ll 1$. Therefore, the eigenvalues of the Jacobian matrix ($J_{EE}$) have negative real parts, and therefore the EE (when it exists) is locally asymptotically stable. We now use the Dulac criterion to establish that no limit cycles exist in $\Omega$. Consider the Dulac function: $\Phi(S, I) = 1/(\beta S I)$, and let the function $F(S, I)$ represent the right hand side of *[1a]*, and let the function $G(S, I)$ represent the right hand side of *[1b]*. Then we have:

$$\frac{\partial(\Phi F)}{\partial S} + \frac{\partial(\Phi G)}{\partial I} = -\frac{(b_R)I}{\beta S^2 I} + q\frac{I^2 - S^2}{\beta S^2 I} < 0 \qquad [7]$$

since $0 < q_R \ll 1$. Thus, system *[1]* has no limit cycles present in $\Omega$. Therefore, the EE is globally asymptotically stable in $\Omega$ when $R_0 > 1$.

### S2 Text. Invasion of a mutant parasite under SI assumptions

Next, we aim to establish under what conditions a mutant pathogen can invade this system. We accomplish this task by first re-writing system *[1]* to include hosts infected with a resident wild-type pathogen, $I_1$, or a novel mutant pathogen, $I_2$ where now, $N = S + I_1 + I_2$:

$$\frac{dS}{dt} = N(b_R - q_R N) - \beta_{r_{1R}} S I_1 - \beta_{r_{2R}} S I_2 - \mu_R S \qquad [8a]$$
$$\frac{dI_1}{dt} = \beta_{r_{1R}} S I_1 - \mu_R I_1 - \alpha_{r_{1R}} I_1 \qquad [8b]$$
$$\frac{dI_2}{dt} = \beta_{r_{2R}} S I_2 - \mu_R I_2 - \alpha_{r_{2R}} I_2 \qquad [8c]$$

Notice that the novel mutant pathogen has a different intrinsic viral growth rate than that of the resident (wild-type) pathogen ($r_{1R} \neq r_{2R}$). It then follows from *[1]* that there exists a mutant-free equilibrium (MFE) in system *[8]* which recapitulates the endemic equilibrium in system *[1]*. To evaluate the local stability of the MFE we can again construct the Jacobian matrix of the new system and evaluate it at the MFE (Note: at the MFE, $I_2^* = 0$ while $S^*$and $I_1^*$ replicate the endemic equilibrium values in *[2a]* and *[2b]*):

$$J_{MFE} = \begin{bmatrix} b_R - 2q_R S^* - 2q_R I_1^* - \beta_{r_{1R}} I_1^* - \mu_R & b_R - 2q_R S^* - 2q_R I_1^* - \beta_{r_{1R}} S^* & b_R - 2q_R S^* - 2q_R I_1^* - \beta_{r_{2R}} S^* \\ \beta_{r_{1R}} I_1^* & \beta_{r_{1R}} S^* - \mu_R - \alpha_{r_{1R}} & 0 \\ 0 & 0 & \beta_{r_{2R}} S^* - \mu_R - \alpha_{r_{2R}} \end{bmatrix} \quad [9]$$

The upper left four quadrats of this matrix exactly replicate the 2x2 Jacobian (*[3a]*) evaluated at the EE ($J_{EE}$), which we now refer to as the resident equilibrium. $J_{MFE}$ is a block-triangular matrix, which we can be broken down into component parts:

$$J_{MFE} = \begin{bmatrix} J_{res} & * \\ 0 & J_{mut} \end{bmatrix} \quad [10]$$

Stability of the MFE requires that the real parts of all eigenvalues of $J_{MFE}$ be negative. Since the matrix is block diagonal, the eigenvalues of $J_{MFE}$ are given by the eigenvalues of the two matrices on the diagonal, $J_{res}$ and $J_{mut}$. We already established that the eigenvalues of $J_{res}$ have negative real parts when $R_0 > 1$. The local stability of the MFE then depends on the sign of the eigenvalue of $J_{mut}$, where $J_{mut} = \beta_2 S_{EE}^* - \mu_R - \alpha_{r_{2R}}$. $J_{mut}$ thus corresponds to the growth rate of a rare mutant in an environment with an endemic resident strain. If the mutant pathogen is able to grow within the population, then:

$$\beta_{r_{2R}} \frac{\mu_R + \alpha_{r_{1R}}}{\beta_{r_{1R}}} - \mu_R - \alpha_{r_{2R}} > 0 \quad [11]$$

or:

$$\frac{\beta_{r_{2R}}}{\mu_R + \alpha_{r_{2R}}} > \frac{\beta_{r_{1R}}}{\mu_R + \alpha_{r_{1R}}} \quad [12]$$

It is evident that the MFE is unstable (allowing for invasion of the mutant strain) when the ratio of transmission over infection duration is larger for the (invading) mutant strain than the resident strain. In general, a pathogen will evolve to maximize this expression, which is known as the invasion fitness, and is given by:

$$\frac{\beta_{r_R}}{\mu_R + \alpha_{r_R}} \quad [13]$$

### S3 Text. Within-host dynamics under SI assumptions

Now that we have determined the invasion fitness for a mutant virus, we can write the population-level rates dependent on within-host dynamics in their within-host terms (transmission: $\beta_{r_{1R}/r_{2R}}$ and virulence: $\alpha_{r_{1R}/r_{2R}}$). This will allow us to derive a selection gradient on the viral growth rate, an intrinsic virus property likely to be carried from one host to another. To do this, we establish a simple within-host model that represents the dynamics of infection *within* each $I_1$ and $I_2$ host as outlined above. We follow Alizon and Van Baalen (2005) (2) to adapt a class of Lotka-Volterra predator-prey-like within-host models (reviewed in (3)), which overcomes some of the constraints of basic predator-prey models of the immune system, chiefly *(i)* by allowing leukocytes to circulate in the absence of infection and *(ii)* by scaling leukocyte growth with virion density, independent of direct leukocyte-virion contact (2,4,5) . This model can be captured by the following system of ODEs, which demonstrates interactions between the virus population ($V_R$) and the leukocyte population ($L_R$) within each infected reservoir host:

$$\frac{dV_R}{dt} = r_R V_R - c_R V_R L_R \quad [14a]$$
$$\frac{dL_R}{dt} = g_{0R} + g_R r_R V_R - m_R L_R \quad [14b]$$

where $r_R$ corresponds to the intrinsic virus growth rate, $c_R$ corresponds to the attack efficacy of the immune system upon contact with the virus, and $g_R$ signifies the recruitment rate of immune cells scaled to the virus growth rate. The parameter, $g_{0R}$, describes the constitutive, baseline leukocyte recruitment in the absence of infection, and $m_R$ gives the natural leukocyte death rate.

This system has two equilibria: a virus free equilibrium at $V_R^* = 0$ and $L_R^* = \frac{g_{0R}}{m_R}$ and an endemic virus equilibrium at $V_R^* = \frac{m_R r_R - c_R g_{0R}}{g_R c_R r_R}$ and $L_R^* = \frac{r_R}{c_R}$.

Building from above, we can then rewrite the component terms of the population-level transmission and virulence rates dependent on within-host dynamics ($\beta_{r_{1R}/r_{2R}}$ and $\alpha_{r_{1R}/r_{2R}}$) in terms of their within-host components, assuming that the within host dynamics are fast relative to the population-level dynamics and converge to the endemic equilibrium (i.e. $m_R r_R > c_R g_{0R}$).

In line with previous work, we assume transmission to be a linear function of viral load (4), which we represent as:

$$\beta_{r_R} = \zeta V_R^*. \quad \textit{[15]}$$

We can also assume that:

$$\alpha_{r_R} = \frac{v r_R V_R^*}{T_{vR}} + \frac{w g_R r_R V_R^*}{T_{wR}}, \quad \textit{[16]}$$

by which infection-induced host mortality ('virulence') results from both virus-induced pathology, a function of the intrinsic virulence of the parasite, $v$, and the parasite growth rate $r_R V_R^*$, and immunopathology, which we model as proportional to the leukocyte growth rate in response to virus expansion, multiplied by the parasite's intrinsic propensity to elicit an inflammatory immune response, given here as $w$. The terms $T_{vR}$ and $T_{wR}$ correspond to host tolerance of virus-induced pathology and immunopathology, respectively.

Expression *[16]* assumes a form of "constant" tolerance, by which both virus pathology and immunopathology are reduced by a constant proportion across the course of infection. For constant tolerance, we assume that $T_{vR} > 1$ and $T_{wR} > 1$. After Miller et al. 2006 (1), we can also model these relationships assuming "complete tolerance" *[17]*, whereby virus pathology and immunopathology are completely eliminated up to a threshold value, beyond which pathology scales proportionally with virus and immune cell growth:

$$\alpha_{r_R} = (v - T_{vR}) r_R V_R^* + (w - T_{wR}) g_R r_R V_R^* \quad \textit{[17]}$$

For complete tolerance, we assume that $0 < T_{vR} < v$ and $0 < T_{wR} < w$.
We consider both constant and complete forms of tolerance in all subsequent analyses. It is important to note that the predictions of the complete form of tolerance are not dependent on the assumption of complete elimination: a more gradual function shows equivalent results (1).

Now, we can rewrite the above equations for transmission (*[15]*) and virulence (*[16]/[17]*) in purely within-host terms, here for constant tolerance:

$$\beta_{r_R} = \frac{\zeta(m_R r_R - c_R g_{0R})}{c_R g_R r_R} \quad \textit{[18]}$$

$$\alpha_{r_R} = \frac{v(m_R r_R - c_R g_{0R})}{c_R g_R T_{vR}} + \frac{w(m_R r_R - c_R g_{0R})}{c_R T_{wR}} \quad \textit{[19]}$$

And here for complete tolerance (note that only the equation for virulence—but not transmission—differs under differing forms of tolerance):

$$\alpha_{r_R} = \frac{(m_R r_R - c_R g_{0R})(v - T_{vR})}{c_R g_R} + \frac{(m_R r_R - c_R g_{0R})(w - T_{wR})}{c_R} \quad \textit{[20]}$$

We can then allow the virus to evolve its within-host growth rate ($r_R$) to maximize its reproductive success (*[13]* for the SI model above). In within-host terms, the invasion fitness takes on the following general form, under assumptions of constant tolerance:

$$\frac{\zeta T_{vR} T_{wR}(m_R r_R - c_R g_{0R})}{r_R T_{wR}(c_R g_R \mu_R T_{vR} - c_R g_{0R} v + m_R r_R v) + g_R r_R w T_{vR}(m_R r_R - c_R g_{0R})} \qquad [21]$$

And under assumptions of complete tolerance:

$$\frac{\zeta(c_R g_{0R} - m_R r_R)}{r_R[m_R r_R(T_{vR} - v + g_R T_{wR} - g_R w) - c_R(g_{0R}(v - T_{vR}) + g_R(\mu_R + g_{0R} T_{wR} - g_{0R} w))]} \qquad [22]$$

We can then determine the selection gradient on $r$ by differentiating *[21]* and *[22]* with respect to $r_R$. This gives us the following expression under assumptions of constant tolerance (differentiating *[21]*):

$$-\frac{\zeta T_{vR} T_{wR}[g_R w T_{vR}(c_R g_{0R} - m_R r_R)^2 + v T_{wR}(c_R g_{0R} - m_R r_R)^2 - c_R^2 g_R g_{0R}^2 \mu_R T_{vR} T_{wR}]}{[r_R T_{wR}(c_R g_R \mu_R T_{vR} - c_R g_{0R} v + m_R r_R v) + g_R r_R w T_{vR}(m_R r_R - c_R g_{0R})]^2} \qquad [23]$$

And assuming complete tolerance (differentiating *[22]*):

$$\frac{\zeta[m_R^2 r_R^2(T_{vR} - v + g_R T_{wR} - g_R w) + 2 c_R g_{0R} m_R r_R(v - T_{vR} + g_R w - g_R T_{wR}) + c_R^2 g_{0R}(g_{0R}(T_{vR} - v) + g_R(\mu + g_{0R} T_{wR} - g_{0R} w))]}{r_R^2[c_R g_{0R}(T_{vR} - v) + c_R g_R(\mu_R + g_{0R}(w - T_{wR})) + m_R r_R(v - T_{vR} + g_R(w - T_{wR}))]^2} \qquad [24]$$

We then set these derivatives equal to 0 and solve for *r*, which we now refer to as $r^*$. The parameter $r^*$ corresponds to the optimal within-host virus growth rate, here under assumptions of constant tolerance:

$$r_R^* = \frac{c_R g_{0R}}{m_R} + \frac{\sqrt{c_R^2 m_R^2 g_R g_{0R} \mu_R T_{vR} T_{wR}(v T_{wR} + g_R w T_{vR})}}{m_R^2(v T_{wR} + g_R w T_{vR})} \qquad [25]$$

And here under assumptions of complete tolerance:

$$r_R^* = \frac{c_R g_{0R}}{m_R} + \frac{c_R^2 g_R g_{0R} \mu_R}{\sqrt{c_R^2 m_R^2 g_R g_{0R} \mu_R (v - T_{vR} + g_R w - g_R T_{wR})}} \qquad [26]$$

We can check that this optimum is a locally stable maximum by proving that the second derivative of the invasion fitness (*[21]* and *[22]*) with respect to $r_R$ is negative at the respective values for $r_R^*$ derived in *[25]* and *[26]*. When we do this, we find that this is, indeed, true, under assumptions of both constant and complete tolerance. This means that *[25]* and *[26]* represent true evolutionarily stable strategies (ESS). Furthermore, we can compute a pairwise invasibility plot to visually demonstrate that the derived expression for $r_R^*$ represents an *ESS* (S1 Fig).

In S1 Fig, we see that all points along the vertical line through the point of intersection fall within the unshaded regions, corresponding to the conditions for a locally stable ESS (6). Additionally, we see that higher degrees of tolerance shift $r_R^*$ higher under both constant and complete tolerance assumptions.

Finally, we can express *[16]* and *[17]* in within-host terms to calculate the virulence which a virus incurs on the reservoir host when evolved to its optimal within-host virus growth rate ($r_R^*$). When we do this, we find that virus-induced virulence at $r_R^*$ (which we call $\alpha_{r_R}^*$) can be expressed as the following, under assumptions of constant tolerance:

$$\alpha_{r_R}^* = \frac{c_R g_{0R} m_R \mu_R (v T_{wR} + g_R w T_{vR})}{\sqrt{c_R^2 m_R^2 g_R g_{0R} \mu_R T_{vR} T_{wR}(v T_{wR} + g_R w T_{vR})}} \qquad [27]$$

And the following, under assumptions of complete tolerance:

$$\alpha^*_{r_R} = \frac{\sqrt{c_R^2 m_R^2 g_R g_{0R} \mu_R (v - T_{vR} + g_R w - g_R T_{wR})}}{c_R g_R m_R} \quad [28]$$

There are several important insights to come out of these simple expressions for optimal viral growth $r_R^*$ and corresponding virulence $\alpha^*_{r_R}$ in a reservoir host. We highlight the relationships between $r_R^*$ and $\alpha^*_{r_R}$ and specific within-host parameters under assumptions of constant tolerance in Fig 2 (main text) and under assumptions of complete tolerance in S2 Fig.

### S4 Text. Spillover virulence in the secondary host

Our primary interest in understanding the evolution of the optimal virus growth rate in a reservoir host is to predict the virulence that such a virus is likely to cause upon spillover to a novel, particularly human, host. Contrary to its established persistent infection in the reservoir host, we assume that the virus will produce an acute infection in the spillover host. To model the dynamics of this acute spillover, we can borrow from Gilchrist and Sasaki 2002 (7), who developed a within-host parasite-leukocyte model in which an immortal leukocyte successfully eradicates the parasite population to near-zero. Here, we modify their acute model to make more comparable to our chronic infection model (*[14a]/[14b]*) and to reflect our differing notation for within-host dynamics.

$$\frac{dV_S}{d\tau} = r_R^* V_S - c_S V_S L_S \quad [29a]$$
$$\frac{dL_S}{d\tau} = r_R^* g_S V_S \quad [29b]$$

where $r_R^*$ represents the reservoir-evolved virus growth rate, but all other terms reflect the within-host conditions of the spillover host: $V_S$ and $L_S$ correspond, respectively, to the spillover host virus and leukocyte populations, $c_S$ is the virus consumption rate upon contact with leukocytes in the spillover host, and $g_S$ is the growth rate of the spillover host leukocyte population in response to virus. Here, the model is expressed in units of $\tau$, which we assume to be short in comparison to the $t$ time units of dynamics at play in the reservoir host population. Rather than solving for virus and leukocyte populations at equilibrium (as in the reservoir host population), we followed Gilchrist and Sasaki 2002 (7) to instead seek to derive an expression for the virus population at the peak of infection ($V_{S_{max}}$), where we anticipate maximum pathology for the spillover host. We then extend this prior work to generate an expression for the average viral load ($V_{S_{avg}}$) in the spillover host, which we use in comparisons of spillover host virulence with reported case fatality rates for zoonoses in the literature.

If we divide equation *[29a]* by equation *[29b]* we can derive a simple time-independent relationship between virus and leukocyte density:

$$\frac{dV_S}{dL_S} = \frac{1}{g_S}\left(1 - \frac{c_S L_S}{r_R^*}\right) \quad [30]$$

If we then let $L_S(0) = 1, V_S(0) = 1$, assuming that both virus and leukocyte populations will be small at $\tau = 0$, we can integrate *[30]* to establish the following relationship between virus and leukocyte:

$$V_S = \frac{1}{g_S} L_S - \frac{c_S}{2 r_R^* g_S}(L_S)^2 + 1 - \frac{1}{g_S} + \frac{c_S}{2 r_S^* g_S} \quad [31]$$

Notice that the above *[31]* is simply a quadratic equation (see *[32]* below):

$$V_S = -\frac{c_S}{2 r_R^* g_S}(L_S)^2 + \frac{1}{g_S} L_S + \left(1 - \frac{1}{g_S} + \frac{c_S}{2 r_S^* g_S}\right) \quad [32]$$

Now, we can take the derivative of equation *[32]* to formulate an expression for $V_{S_{max}}$, the maximum viral load, which should precede the end of acute infection and the point of host recovery at the maximum duration of infection:

$$V_{S_{max}} = \frac{r_R^*}{g_S c_S} - \frac{r_R^*}{2 g_S c_S} + 1 - \frac{1}{g_S} + \frac{c_S}{2 r_R^* g_S} \qquad [33]$$

Now, extending previous work (7), we can calculate the average viral load by taking the integral of equation *[32]* from $V_S(0)$ to $V_{S_{max}}$ and dividing by the duration of that interval. From this exercise, we can express the average value of equation *[32]* as:

$$V_{S_{avg}} = \frac{r_R^* \sqrt{c_S^2 + 2 r_R^* (g_S - 1) c + \left(r_R^*\right)^2} + (r_R^*)^2 + 4 r_R^* (g-1) c + 2 c_S^2}{6 c_S g_S r_R^*} \qquad [34]$$

Now, with this established expression for $V_{S_{avg}}$, we can adapt equations *[16]* and *[17]* to reflect the acute within-host dynamics of "spillover virulence", which we model (as before) as a combination of mortality induced from direct virus pathology and from immunopathology. As in the reservoir host system, we can also model the mitigating impact of tolerance on the two mechanisms of virulence, here under assumptions of constant tolerance:

$$\alpha_S = V_{S_{avg}} \left( \frac{r_R^* v}{T_{vS}} + \frac{g_S w r_R^*}{T_{wS}} \right) \qquad [35]$$

In our expression for spillover virulence, we model the growth rate of the virus at its evolutionary optimum evolved in the reservoir host ($r_R^*$), and we represent a virus's intrinsic virulence ($v$) and propensity to elicit an inflammatory immune response ($w$) as unchanged from one host to the next. Inspired by findings in the literature that report higher virulence in cross-species infections between hosts separated by larger phylogenetic distances (8–12), we model spillover host tolerance of virus-induced pathology ($T_{vS}$) as a decreasing function of increasing phylogenetic distance between the reservoir and secondary host. All other immune-related parameters take on characteristics of the spillover host: $g_S$ is the spillover host's leukocyte growth rate, and $T_{wS}$ corresponds to the spillover host's tolerance of immunopathology.

We can also represent spillover virulence under assumptions of complete tolerance:

$$\alpha_S = V_{S_{avg}} \left( r_R^* (v - T_{vS}) + r_R^* g_{SR} (w - T_{wS}) \right) \qquad [36]$$

**Table S1. Order-specific parameter values for within-host nested model.***

| | $\mu_R$, reservoir host background mortality (days$^{-1}$) | $T_{wR}$, reservoir host tolerance of immunopathology *(constant)* | $T_{wR}$, reservoir host tolerance of mmunopatholog *(complete)* | $g_{0R}$, reservoir host magnitude of constitutive immunity (days$^{-1}$) | $T_{vS}$, human tolerance of viral pathology *(constant)* | $T_{vS}$, human tolerance of viral pathology *(complete)* | $r_R^*$, optimal virus growth rate (days$^{-1}$) *(constant)* | $r_R^*$, optimal virus growth rate (days$^{-1}$) *(complete)* | $\alpha_S$, relative spillover virulence (days$^{-1}$) *(constant)* | $\alpha_S$, relative spillover virulence (days$^{-1}$) *(complete)* |
|---|---|---|---|---|---|---|---|---|---|---|
| Afrosoricida | 8.58x10$^4$ [6.56x10$^4$-1.06x10$^3$] | 1.42 [1.31-1.53] | 0.42 [0.305-0.535] | 0.0613 [0.0428-0.0798] | 1.45 | 0.451 | 0.7 [0.49-0.911] | 0.703 [0.491-0.915] | 0.328 [0.168-0.615] | 0.293 [0.152-0.553] |
| Carnivora | 1.75x10$^4$ [1.26x10$^4$-2.24x10$^4$] | 1.64 [1.61-1.67] | 0.638 [0.609-0.667] | 0.0484 [0.0439-0.0529] | 1.48 | 0.478 | 0.531 [0.48-0.583] | 0.533 [0.481-0.585] | 0.192 [0.159-0.341] | 0.17 [0.141-0.303] |
| Cetartiodactyla | 1.54x10$^4$ [1.09x10$^4$-2.00x10$^4$] | 1.49 [1.46-1.52] | 0.491 [0.464-0.518] | 0.0283 [0.021-0.0357] | 1.48 | 0.478 | 0.314 [0.232-0.396] | 0.315 [0.233-0.397] | 0.0764 [0.0478-0.243] | 0.0696 [0.0447-0.215] |
| Chiroptera | 2.54x10$^4$ [1.83x10$^4$-3.24x10$^4$] | 2.01 [1.96-2.05] | 1.01 [0.964-1.05] | 0.0736 [0.0644-0.0828] | 1.48 | 0.478 | 0.808 [0.704-0.912] | 0.816 [0.71-0.922] | 0.428 [0.327-0.611] | 0.38 [0.29-0.549] |
| Cingulata | 1.64x10$^4$ [0-3.90x10$^4$] | 1.74 [1.61-1.86] | 0.737 [0.61-0.864] | 0.0394 [0.0254-0.0533] | 1.45 | 0.451 | 0.434 [0.267-0.597] | 0.436 [0-0.603] | 0.135 [0.0609-0.353] | 0.124 [0-0.319] |
| Dasyuromorphia | 8.21x10$^4$ [7.20x10$^4$-9.22x10$^4$] | 1.4 [1.34-1.46] | 0.399 [0.34-0.458] | 0.0293 [0.0136-0.0451] | 1.11 | 0.111 | 0.346 [0.167-0.524] | 0.347 [0.167-0.526] | 0.127 [0.0587-0.349] | 0.14 [0.0744-0.341] |
| Dermoptera | 1.57x10$^4$ [0-6.09x10$^4$] | 1.73 [1.51-1.95] | 0.731 [0.509-0.953] | --- | --- | --- | --- | --- | --- | --- |
| Didelphimorphia | 9.41x10$^4$ [7.81x10$^4$-1.10x10$^3$] | 1.27 [1.17-1.36] | 0.265 [0.173-0.357] | 0.0528 [0.0293-0.0763] | 1.11 | 0.111 | 0.608 [0.343-0.872] | 0.609 [0.343-0.874] | 0.306 [0.125-0.652] | 0.31 [0.138-0.637] |
| Diprotodontia | 2.97x10$^4$ [2.18x10$^4$-3.75x10$^4$] | 1.55 [1.51-1.6] | 0.554 [0.507-0.6] | 0.0288 [0.0206-0.0369] | 1.11 | 0.111 | 0.325 [0.233-0.417] | 0.327 [0.234-0.419] | 0.117 [0.0794-0.288] | 0.13 [0.0942-0.281] |
| Eulipotyphla | 1.10x10$^3$ [1.01x10$^3$-1.19x10$^3$] | 1.36 [1.3-1.41] | 0.359 [0.304-0.415] | 0.0648 [0.0413-0.0883] | 1.48 | 0.478 | 0.746 [0.483-1.01] | 0.748 [0.484-1.01] | 0.366 [0.161-0.713] | 0.321 [0.143-0.633] |
| Hyracoidea | 2.21x10$^4$ [0-5.41x10$^4$] | 1.58 [1.41-1.75] | 0.58 [0.409-0.751] | 0.0619 [0.0384-0.0855] | 1.45 | 0.451 | 0.68 [0.403-0.952] | 0.682 [0-0.959] | 0.31 [0.119-0.658] | 0.277 [0-0.594] |
| Lagomorpha | 4.04x10$^4$ [2.33x10$^4$-5.75x10$^4$] | 1.48 [1.39-1.58] | 0.485 [0.387-0.583] | --- | 1.52 | 0.516 | --- | --- | --- | --- |
| Macroscelidea | 7.87x10$^4$ [5.26x10$^4$-1.05x10$^3$] | 1.5 [1.35-1.64] | 0.496 [0.352-0.64] | --- | 1.45 | 0.451 | --- | --- | --- | --- |
| Microbiotheria | 8.56x10$^4$ [2.16x10$^4$-1.50x10$^3$] | 1.48 [1.21-1.76] | 0.481 [0.208-0.755] | --- | 1.11 | 0.111 | --- | --- | --- | --- |
| Monotremata | 8.88x10$^5$ [0-4.58x10$^4$] | 1.88 [1.69-2.07] | 0.882 [0.691-1.07] | 0.0815 [0.063-0.1] | 1 | 0 | 0.878 [0.661-1.11] | 0.881 [0-1.12] | 0.643 [0.384-1] | 0.639 [0-1] |
| Notoryctemorphia | 1.83x10$^3$ [1.19x10$^3$-2.47x10$^3$] | 1.27 [1-1.55] | 0.274 [0-0.548] | --- | 1.11 | 0.111 | --- | --- | --- | --- |
| Peramelemorphia | 5.83x10$^4$ [3.41x10$^4$-8.24x10$^4$] | 1.31 [1.18-1.45] | 0.311 [0.176-0.446] | 0.0603 [0.0417-0.0788] | 1.11 | 0.111 | 0.678 [0.467-0.89] | 0.679 [0.467-0.893] | 0.372 [0.197-0.672] | 0.373 [0.206-0.658] |
| Perissodactyla | 7.74x10$^5$ [0-2.48x10$^4$] | 1.61 [1.51-1.7] | 0.606 [0.508-0.704] | 0.0331 [0.00962-0.0567] | 1.48 | 0.478 | 0.361 [0.101-0.625] | 0.362 [0-0.628] | 0.0963 [0.0166-0.369] | 0.0868 [0-0.328] |

| | | | | | | | | | | |
|---|---|---|---|---|---|---|---|---|---|---|
| Pholidota | 3.17x10$^{4}$ [0-6.37x10$^{4}$] | 1.45 [1.28-1.62] | 0.453 [0.282-0.624] | --- | 1.48 | 0.478 | --- | --- | --- | --- |
| Pilosa | 3.11x10$^{4}$ [6.93x10$^{5}$-5.53x10$^{4}$] | 1.59 [1.46-1.73] | 0.592 [0.457-0.727] | 0.0477 [0.0319-0.0635] | 1.45 | 0.451 | 0.532 [0.347-0.714] | 0.534 [0.348-0.719] | 0.195 [0.0923-0.437] | 0.176 [0.0857-0.394] |
| Primates | 1.20x10$^{4}$ [6.27x10$^{5}$-1.77x10$^{4}$] | 1.89 [1.85-1.92] | 0.888 [0.855-0.922] | 0.0545 [0.0482-0.0608] | 2 | 1 | 0.593 [0.52-0.665] | 0.597 [0.522-0.67] | 0.19 [0.144-0.352] | 0.0791 [0.0504-0.224] |
| Proboscidea | 3.70x10$^{5}$ [0-4.06x10$^{4}$] | 1.69 [1.5-1.89] | 0.694 [0.503-0.886] | 0.0553 [0.0318-0.0789] | 1.45 | 0.451 | 0.593 [0.334-0.875] | 0.594 [0-0.882] | 0.238 [0.0868-0.578] | 0.214 [0-0.523] |
| Rodentia | 6.21x10$^{4}$ [5.79x10$^{4}$-6.64x10$^{4}$] | 1.5 [1.47-1.52] | 0.496 [0.47-0.521] | 0.0452 [0.0376-0.0528] | 1.52 | 0.516 | 0.516 [0.431-0.601] | 0.518 [0.433-0.603] | 0.178 [0.128-0.349] | 0.153 [0.11-0.305] |
| Scandentia | 3.92x10$^{4}$ [1.06x10$^{4}$-6.78x10$^{4}$] | 1.66 [1.51-1.82] | 0.662 [0.507-0.818] | 0.0185 [1x107-0.037] | 1.53 | 0.529 | 0.216 [2.68x105-0.43] | 0.218 [2.81x105-0.435 | 0.04 [0-0.254] | 0.0343 [0-0.222] |
| Sirenia | 7.39x10$^{5}$ [0-4.43x10$^{4}$] | 1.64 [1.45-1.83] | 0.64 [0.449-0.832] | --- | 1.45 | 0.451 | --- | --- | --- | --- |
| Tubulidentata | 1.02x10$^{4}$ [0-7.42x10$^{4}$] | 1.61 [1.34-1.89] | 0.614 [0.34-0.888] | 0.0382 [0.0147-0.0618] | 1.45 | 0.451 | 0.417 [0.155-0.704] | 0.419 [0-0.713] | 0.126 [0.029-0.429] | 0.116 [0-0.39] |

*Derived parameter values (depicted as *mean [lower confidence interval – upper confidence interval]*) computed from linear regression and mixed effects models are shown across all 26 mammalian orders ($\mu_R, g_{0R}, T_{wR}, T_{vS}$), in addition to the $r_R^*$ and $\alpha_S$ values computed from their inputs. Confidence intervals are 95% derived via standard error. All other parameters involved in the calculation of $r_R^*$ and $\alpha_S$ are listed in main text Table 1.

**S2 Table. Model summary outputs for within-host parameter estimation.***

**A.** Summary of linear regression model to predict $\boldsymbol{\mu_R}$

**Formula:** *lm(mortality_rate~order, data = pan.dat);* **Number observations:** 1060; $\mathbf{R^2}$=0.42

| Term | Estimate | Lci-Uci | Statistic | P-value |
|---|---|---|---|---|
| Intercept (Diprotodontia) | 0.11 | 0.08 - 0.14 | T: 7.38 | <0.001*** |
| Rodentia | 0.12 | 0.09 - 0.15 | T: 7.1 | <0.001*** |
| Carnivora | -0.04 | 0-0.07 | T: -2.58 | 0.01** |
| Cetartiodactyla | -0.05 | 0-0.07 | T: -3.07 | <0.001*** |
| Primates | -0.06 | 0-0.07 | T: -3.56 | <0.001*** |
| Afrosoricida | 0.2 | 0.13 - 0.28 | T: 5.07 | <0.001*** |
| Chiroptera | -0.02 | 0-0.07 | T: -0.8 | 0.42 |
| Dasyuromorphia | 0.19 | 0.14 - 0.24 | T: 8.02 | <0.001*** |
| Eulipotyphla | 0.29 | 0.25 - 0.34 | T: 12.82 | <0.001*** |
| Pilosa | 0.01 | 0-0.19 | T: 0.11 | 0.91 |
| Didelphimorphia | 0.24 | 0.17 - 0.3 | T: 7.09 | <0.001*** |
| Cingulata | -0.05 | 0-0.18 | T: -1.09 | 0.28 |
| Perissodactyla | -0.08 | 0-0.14 | T: -2.28 | 0.02* |
| Dermoptera | -0.05 | 0-0.34 | T: -0.6 | 0.55 |
| Hyracoidea | -0.03 | 0-0.24 | T: -0.45 | 0.65 |
| Microbiotheria | 0.2 | 0-0.47 | T: 1.7 | 0.09 |
| Sirenia | -0.08 | 0-0.28 | T: -1.16 | 0.25 |
| Peramelemorphia | 0.1 | 0.01 - 0.2 | T: 2.2 | 0.03 |
| Macroscelidea | 0.18 | 0.08 - 0.28 | T: 3.53 | <0.001*** |
| Proboscidea | -0.09 | 0-0.27 | T: -1.35 | 0.18 |
| Lagomorpha | 0.04 | 0-0.14 | T: 1.12 | 0.26 |
| Pholidota | 0.01 | 0-0.24 | T: 0.12 | 0.9 |
| Notoryctemorphia | 0.56 | 0.32 - 0.79 | T: 4.65 | <0.001*** |
| Monotremata | -0.08 | 0-0.27 | T: -1.08 | 0.28 |
| Tubulidentata | -0.07 | 0-0.47 | T: -0.59 | 0.55 |
| Scandentia | 0.03 | 0-0.21 | T: 0.63 | 0.53 |

**B.** Summary of linear mixed effect regression model to estimate $\boldsymbol{T_{wR}}$

**Formula:** *lmer(log10_max_lifespan_yrs ~ log10mass_g + (1 | order), data = pan.dat)*
**Number observations:** 1060; **Groups:** 26

| Term | Estimate | Lci-Uci | Statistic | P-value |
|---|---|---|---|---|
| (Intercept) | 0.392 | 0.295-0.488 | T: 7.96 | <0.001*** |
| log10mass (grams) | .199 | 0.185-0.214 | T: 26.55 | <0.001*** |
| order (random effect) | Variance: 0.041 | Std Dev: 0.202 | | |

**C.** Summary of linear mixed effect regression model to estimate $\boldsymbol{g_0}$

**Formula:** *lmer(log10_neutrophil_count ~ log10mass_g + BMR_W + (1 | order), data = pan.dat)*
**Number observations:** 144; **Groups**: 19

| Term | Estimate | Lci-Uci | Statistic | P-value |
|---|---|---|---|---|
| (Intercept) | -0.245 | -0.422 - -068 | T: 2.72 | 0.008** |
| log10mass (grams) | 0.212 | 0.168-0.255 | T: 9.53 | <0.001*** |
| BMR (W) | -0.000368 | -0.000103- - 0.000633 | T: -2.72 | 0.01* |
| order (random effect) | Variance: 0.041 | Std Dev: 0.203 | | |

**D**. Summary of generalized additive model to summarize $\boldsymbol{\alpha_S}$ by order from literature

**Formula:** *alpha ~ s(vFamily, bs = "re") + s(hOrder, bs = "re") + s(VirusPubs, k = 7, bs = "tp") + s(spill_type, bs = "re") + s(IsVectorBorne, bs = "re");* **Deviance explained:** 88.2%; $\mathbf{R^2}$**:** 0.873; **N** = 75

| Term | Estimate | Lci-Uci | Statistic | P-value | Effective degrees of freedom |
|---|---|---|---|---|---|
| Intercept *smoothing:* | -4.55 | [-6.15 – - 2.95] | *z:* -5.57 | <.001*** | |
| virus family | | | $\chi^2$: 612 | .00387** | 9.50 |
| order | | | $\chi^2$: 1161 | <.001*** | 4.74 |
| virus spp. pub count | | | $\chi^2$: 5.77 | .0163* | 1.00 |
| spillover type | | | $\chi^2$: 67.9 | .00856** | 0.867 |
| vector-borne status | | | $\chi^2$: 98.9 | .0249* | 0.806 |

*Outputs from fitted linear and linear mixed effect regression models used to generate within-host parameter values. Projections for model A and order-level effects in B and C are visualized in main text Fig 3C. Order-level predictions for each parameter are visualized in S4 Fig and presented in S1 Table. Significance at p < 0.001***, 0.01**, 0.05*.

**S3 Table. Sensitivity analysis and comparison of influence of individual parameter estimates on overall fit of nested model to data.**

| Order | Tolerance | Fitted Par | Fit Type* | Par Value (this fit) | Par Value (glm fit) | $\alpha_S$ (this fit) | $\alpha_S$ (glm fit) | $\alpha_S$ (lit) |
|---|---|---|---|---|---|---|---|---|
| Carnivora | constant | $T_{wR}$ | one profiled; rest of parameters constant | 2 | 1.64 | 0.63 | 0.61 | 0.65 |
| Carnivora | constant | $T_{vS}$ | one profiled; rest of parameters constant | 1.39 | 1.48 | 0.65 | 0.61 | 0.65 |
| Carnivora | constant | $g_{0R}$ | one profiled; rest of parameters constant | 0.05 | 0.05 | 0.63 | 0.61 | 0.65 |
| Carnivora | constant | $T_{wR}$ | one profiled; glm fit rest of parameters | 2 | 1.64 | 0.61 | 0.61 | 0.65 |
| Carnivora | constant | $T_{vS}$ | one profiled; glm fit rest of parameters | 1.3 | 1.48 | 0.64 | 0.61 | 0.65 |
| Carnivora | constant | $g_{0R}$ | one profiled; glm fit rest of parameters | 0.05 | 0.05 | 0.64 | 0.61 | 0.65 |
| Cetartiodactyla | constant | $T_{wR}$ | one profiled; rest of parameters constant | 1 | 1.49 | 0.62 | 0.42 | 0.42 |
| Cetartiodactyla | constant | $T_{vS}$ | one profiled; rest of parameters constant | 2 | 1.48 | 0.56 | 0.42 | 0.42 |
| Cetartiodactyla | constant | $g_{0R}$ | one profiled; rest of parameters constant | 0.03 | 0.03 | 0.43 | 0.42 | 0.42 |
| Cetartiodactyla | constant | $T_{wR}$ | one profiled; glm fit rest of parameters | 1.49 | 1.49 | 0.42 | 0.42 | 0.42 |
| Cetartiodactyla | constant | $T_{vS}$ | one profiled; glm fit rest of parameters | 1.47 | 1.48 | 0.42 | 0.42 | 0.42 |
| Cetartiodactyla | constant | $g_{0R}$ | one profiled; glm fit rest of parameters | 0.03 | 0.03 | 0.43 | 0.42 | 0.42 |
| Chiroptera | constant | $T_{wR}$ | one profiled; rest of parameters constant | 2 | 2.01 | 0.63 | 1.01 | 1.01 |
| Chiroptera | constant | $T_{vS}$ | one profiled; rest of parameters constant | 1 | 1.48 | 0.76 | 1.01 | 1.01 |
| Chiroptera | constant | $g_{0R}$ | one profiled; rest of parameters constant | 0.07 | 0.07 | 0.95 | 1.01 | 1.01 |
| Chiroptera | constant | $T_{wR}$ | one profiled; glm fit rest of parameters | 2 | 2.01 | 1.01 | 1.01 | 1.01 |
| Chiroptera | constant | $T_{vS}$ | one profiled; glm fit rest of parameters | 1.47 | 1.48 | 1.01 | 1.01 | 1.01 |
| Chiroptera | constant | $g_{0R}$ | one profiled; glm fit rest of parameters | 0.07 | 0.07 | 0.95 | 1.01 | 1.01 |
| Diprotodontia | constant | $T_{wR}$ | one profiled; rest of parameters constant | 1 | 1.55 | 0.63 | 0.48 | 0.47 |
| Diprotodontia | constant | $T_{vS}$ | one profiled; rest of parameters constant | 2 | 1.11 | 0.57 | 0.48 | 0.47 |
| Diprotodontia | constant | $g_{0R}$ | one profiled; rest of parameters constant | 0.03 | 0.03 | 0.43 | 0.48 | 0.47 |
| Diprotodontia | constant | $T_{wR}$ | one profiled; glm fit rest of parameters | 1 | 1.55 | 0.48 | 0.48 | 0.47 |
| Diprotodontia | constant | $T_{vS}$ | one profiled; glm fit rest of parameters | 1.2 | 1.11 | 0.47 | 0.48 | 0.47 |
| Diprotodontia | constant | $g_{0R}$ | one profiled; glm fit rest of parameters | 0.03 | 0.03 | 0.5 | 0.48 | 0.47 |

| | | | | | | | | |
|---|---|---|---|---|---|---|---|---|
| Eulipotyphla | constant | $T_{wR}$ | one profiled; rest of parameters constant | 1 | 1.36 | 0.66 | 0.9 | 0.54 |
| Eulipotyphla | constant | $T_{vS}$ | one profiled; rest of parameters constant | 2 | 1.48 | 0.6 | 0.9 | 0.54 |
| Eulipotyphla | constant | $g_{0R}$ | one profiled; rest of parameters constant | 0.04 | 0.06 | 0.55 | 0.9 | 0.54 |
| Eulipotyphla | constant | $T_{wR}$ | one profiled; glm fit rest of parameters | 1 | 1.36 | 0.9 | 0.9 | 0.54 |
| Eulipotyphla | constant | $T_{vS}$ | one profiled; glm fit rest of parameters | 2 | 1.48 | 0.8 | 0.9 | 0.54 |
| Eulipotyphla | constant | $g_{0R}$ | one profiled; glm fit rest of parameters | 0.04 | 0.06 | 0.55 | 0.9 | 0.54 |
| Perissodactyla | constant | $T_{wR}$ | one profiled; rest of parameters constant | 1 | 1.61 | 0.61 | 0.45 | 0.55 |
| Perissodactyla | constant | $T_{vS}$ | one profiled; rest of parameters constant | 2 | 1.48 | 0.55 | 0.45 | 0.55 |
| Perissodactyla | constant | $g_{0R}$ | one profiled; rest of parameters constant | 0.04 | 0.03 | 0.51 | 0.45 | 0.55 |
| Perissodactyla | constant | $T_{wR}$ | one profiled; glm fit rest of parameters | 2 | 1.61 | 0.45 | 0.45 | 0.55 |
| Perissodactyla | constant | $T_{vS}$ | one profiled; glm fit rest of parameters | 1 | 1.48 | 0.54 | 0.45 | 0.55 |
| Perissodactyla | constant | $g_{0R}$ | one profiled; glm fit rest of parameters | 0.04 | 0.03 | 0.51 | 0.45 | 0.55 |
| Primates | constant | $T_{wR}$ | one profiled; rest of parameters constant | 1 | 1.89 | 0.62 | 0.61 | 0.46 |
| Primates | constant | $T_{vS}$ | one profiled; rest of parameters constant | 2 | 2 | 0.55 | 0.61 | 0.46 |
| Primates | constant | $g_{0R}$ | one profiled; rest of parameters constant | 0.03 | 0.05 | 0.43 | 0.61 | 0.46 |
| Primates | constant | $T_{wR}$ | one profiled; glm fit rest of parameters | 1 | 1.89 | 0.6 | 0.61 | 0.46 |
| Primates | constant | $T_{vS}$ | one profiled; glm fit rest of parameters | 2 | 2 | 0.61 | 0.61 | 0.46 |
| Primates | constant | $g_{0R}$ | one profiled; glm fit rest of parameters | 0.04 | 0.05 | 0.46 | 0.61 | 0.46 |
| Rodentia | constant | $T_{wR}$ | one profiled; rest of parameters constant | 1 | 1.5 | 0.64 | 0.59 | 0.52 |
| Rodentia | constant | $T_{vS}$ | one profiled; rest of parameters constant | 2 | 1.52 | 0.58 | 0.59 | 0.52 |
| Rodentia | constant | $g_{0R}$ | one profiled; rest of parameters constant | 0.04 | 0.05 | 0.54 | 0.59 | 0.52 |
| Rodentia | constant | $T_{wR}$ | one profiled; glm fit rest of parameters | 1 | 1.5 | 0.58 | 0.59 | 0.52 |
| Rodentia | constant | $T_{vS}$ | one profiled; glm fit rest of parameters | 2 | 1.52 | 0.53 | 0.59 | 0.52 |
| Rodentia | constant | $g_{0R}$ | one profiled; glm fit rest of parameters | 0.04 | 0.05 | 0.53 | 0.59 | 0.52 |
| Carnivora | complete | $T_{wR}$ | one profiled; rest of parameters constant | 1 | 0.64 | 0.56 | 0.55 | 0.59 |
| Carnivora | complete | $T_{vS}$ | one profiled; rest of parameters constant | 0.43 | 0.48 | 0.59 | 0.55 | 0.59 |
| Carnivora | complete | $g_{0R}$ | one profiled; rest of parameters constant | 0.05 | 0.05 | 0.57 | 0.55 | 0.59 |

| | | | | | | | | |
|---|---|---|---|---|---|---|---|---|
| Carnivora | complete | $T_{wR}$ | one profiled; glm fit rest of parameters | 1 | 0.64 | 0.56 | 0.55 | 0.59 |
| Carnivora | complete | $T_{vS}$ | one profiled; glm fit rest of parameters | 0.38 | 0.48 | 0.59 | 0.55 | 0.59 |
| Carnivora | complete | $g_{0R}$ | one profiled; glm fit rest of parameters | 0.05 | 0.05 | 0.58 | 0.55 | 0.59 |
| Cetartiodactyla | complete | $T_{wR}$ | one profiled; rest of parameters constant | 0 | 0.49 | 0.56 | 0.38 | 0.38 |
| Cetartiodactyla | complete | $T_{vS}$ | one profiled; rest of parameters constant | 0.96 | 0.48 | 0.37 | 0.38 | 0.38 |
| Cetartiodactyla | complete | $g_{0R}$ | one profiled; rest of parameters constant | 0.03 | 0.03 | 0.38 | 0.38 | 0.38 |
| Cetartiodactyla | complete | $T_{wR}$ | one profiled; glm fit rest of parameters | 0.49 | 0.49 | 0.38 | 0.38 | 0.38 |
| Cetartiodactyla | complete | $T_{vS}$ | one profiled; glm fit rest of parameters | 0.47 | 0.48 | 0.38 | 0.38 | 0.38 |
| Cetartiodactyla | complete | $g_{0R}$ | one profiled; glm fit rest of parameters | 0.03 | 0.03 | 0.39 | 0.38 | 0.38 |
| Chiroptera | complete | $T_{wR}$ | one profiled; rest of parameters constant | 1 | 1.01 | 0.57 | 0.92 | 0.92 |
| Chiroptera | complete | $T_{vS}$ | one profiled; rest of parameters constant | 0 | 0.48 | 0.77 | 0.92 | 0.92 |
| Chiroptera | complete | $g_{0R}$ | one profiled; rest of parameters constant | 0.07 | 0.07 | 0.85 | 0.92 | 0.92 |
| Chiroptera | complete | $T_{wR}$ | one profiled; glm fit rest of parameters | 1 | 1.01 | 0.92 | 0.92 | 0.92 |
| Chiroptera | complete | $T_{vS}$ | one profiled; glm fit rest of parameters | 0.47 | 0.48 | 0.92 | 0.92 | 0.92 |
| Chiroptera | complete | $g_{0R}$ | one profiled; glm fit rest of parameters | 0.07 | 0.07 | 0.87 | 0.92 | 0.92 |
| Diprotodontia | complete | $T_{wR}$ | one profiled; rest of parameters constant | 0 | 0.55 | 0.56 | 0.48 | 0.42 |
| Diprotodontia | complete | $T_{vS}$ | one profiled; rest of parameters constant | 0.86 | 0.11 | 0.42 | 0.48 | 0.42 |
| Diprotodontia | complete | $g_{0R}$ | one profiled; rest of parameters constant | 0.03 | 0.03 | 0.39 | 0.48 | 0.42 |
| Diprotodontia | complete | $T_{wR}$ | one profiled; glm fit rest of parameters | 0 | 0.55 | 0.48 | 0.48 | 0.42 |
| Diprotodontia | complete | $T_{vS}$ | one profiled; glm fit rest of parameters | 0.33 | 0.11 | 0.42 | 0.48 | 0.42 |
| Diprotodontia | complete | $g_{0R}$ | one profiled; glm fit rest of parameters | 0.02 | 0.03 | 0.42 | 0.48 | 0.42 |
| Eulipotyphla | complete | $T_{wR}$ | one profiled; rest of parameters constant | 0 | 0.36 | 0.59 | 0.82 | 0.49 |
| Eulipotyphla | complete | $T_{vS}$ | one profiled; rest of parameters constant | 0.75 | 0.48 | 0.49 | 0.82 | 0.49 |
| Eulipotyphla | complete | $g_{0R}$ | one profiled; rest of parameters constant | 0.04 | 0.06 | 0.49 | 0.82 | 0.49 |
| Eulipotyphla | complete | $T_{wR}$ | one profiled; glm fit rest of parameters | 0 | 0.36 | 0.81 | 0.82 | 0.49 |
| Eulipotyphla | complete | $T_{vS}$ | one profiled; glm fit rest of parameters | 1 | 0.48 | 0.52 | 0.82 | 0.49 |
| Eulipotyphla | complete | $g_{0R}$ | one profiled; glm fit rest of parameters | 0.04 | 0.06 | 0.5 | 0.82 | 0.49 |

| | | | | | | | | |
|---|---|---|---|---|---|---|---|---|
| Perissodactyla | complete | $T_{wR}$ | one profiled; rest of parameters constant | 0 | 0.61 | 0.55 | 0.41 | 0.5 |
| Perissodactyla | complete | $T_{vS}$ | one profiled; rest of parameters constant | 0.64 | 0.48 | 0.5 | 0.41 | 0.5 |
| Perissodactyla | complete | $g_{0R}$ | one profiled; rest of parameters constant | 0.04 | 0.03 | 0.46 | 0.41 | 0.5 |
| Perissodactyla | complete | $T_{wR}$ | one profiled; glm fit rest of parameters | 1 | 0.61 | 0.41 | 0.41 | 0.5 |
| Perissodactyla | complete | $T_{vS}$ | one profiled; glm fit rest of parameters | 0.16 | 0.48 | 0.5 | 0.41 | 0.5 |
| Perissodactyla | complete | $g_{0R}$ | one profiled; glm fit rest of parameters | 0.04 | 0.03 | 0.47 | 0.41 | 0.5 |
| Primates | complete | $T_{wR}$ | one profiled; rest of parameters constant | 0 | 0.89 | 0.55 | 0.39 | 0.42 |
| Primates | complete | $T_{vS}$ | one profiled; rest of parameters constant | 0.86 | 1 | 0.41 | 0.39 | 0.42 |
| Primates | complete | $g_{0R}$ | one profiled; rest of parameters constant | 0.03 | 0.05 | 0.38 | 0.39 | 0.42 |
| Primates | complete | $T_{wR}$ | one profiled; glm fit rest of parameters | 1 | 0.89 | 0.39 | 0.39 | 0.42 |
| Primates | complete | $T_{vS}$ | one profiled; glm fit rest of parameters | 0.95 | 1 | 0.41 | 0.39 | 0.42 |
| Primates | complete | $g_{0R}$ | one profiled; glm fit rest of parameters | 0.06 | 0.05 | 0.44 | 0.39 | 0.42 |
| Rodentia | complete | $T_{wR}$ | one profiled; rest of parameters constant | 0 | 0.5 | 0.58 | 0.52 | 0.47 |
| Rodentia | complete | $T_{vS}$ | one profiled; rest of parameters constant | 0.77 | 0.52 | 0.47 | 0.52 | 0.47 |
| Rodentia | complete | $g_{0R}$ | one profiled; rest of parameters constant | 0.04 | 0.05 | 0.48 | 0.52 | 0.47 |
| Rodentia | complete | $T_{wR}$ | one profiled; glm fit rest of parameters | 0 | 0.5 | 0.52 | 0.52 | 0.47 |
| Rodentia | complete | $T_{vS}$ | one profiled; glm fit rest of parameters | 0.65 | 0.52 | 0.47 | 0.52 | 0.47 |
| Rodentia | complete | $g_{0R}$ | one profiled; glm fit rest of parameters | 0.04 | 0.05 | 0.47 | 0.52 | 0.47 |

* Two sensitivity analyses were conducted: in the first, all within-host parameters were held constant across all mammalian orders at values listed in Table 1, excepting $\mu_R$which we varied by order according to values presented in S1 Table, and one additional parameter (either $T_{wR}, g_{0R}$, or $T_{vS}$) which we optimized. In these exercises, the parameter under optimization (listed here) was profiled until we recovered the best log-likelihood fit to the case fatality data recovered from the literature for that particular order. In the second exercise, parameter values for $\mu_R$, $T_{wR}$, $g_{0R}$, and $T_{vS}$ were varied across orders at values recovered from linear modeling of publicly available life history data, as presented in S1 Table. Here, we optimized one of three variable parameters (either $T_{wR}, g_{0R}$, or $T_{vS}$) further while holding all other parameters at S1 Table values. Resulting model fits and linear regression of observed vs. predicted values are visualized in S9 Fig.

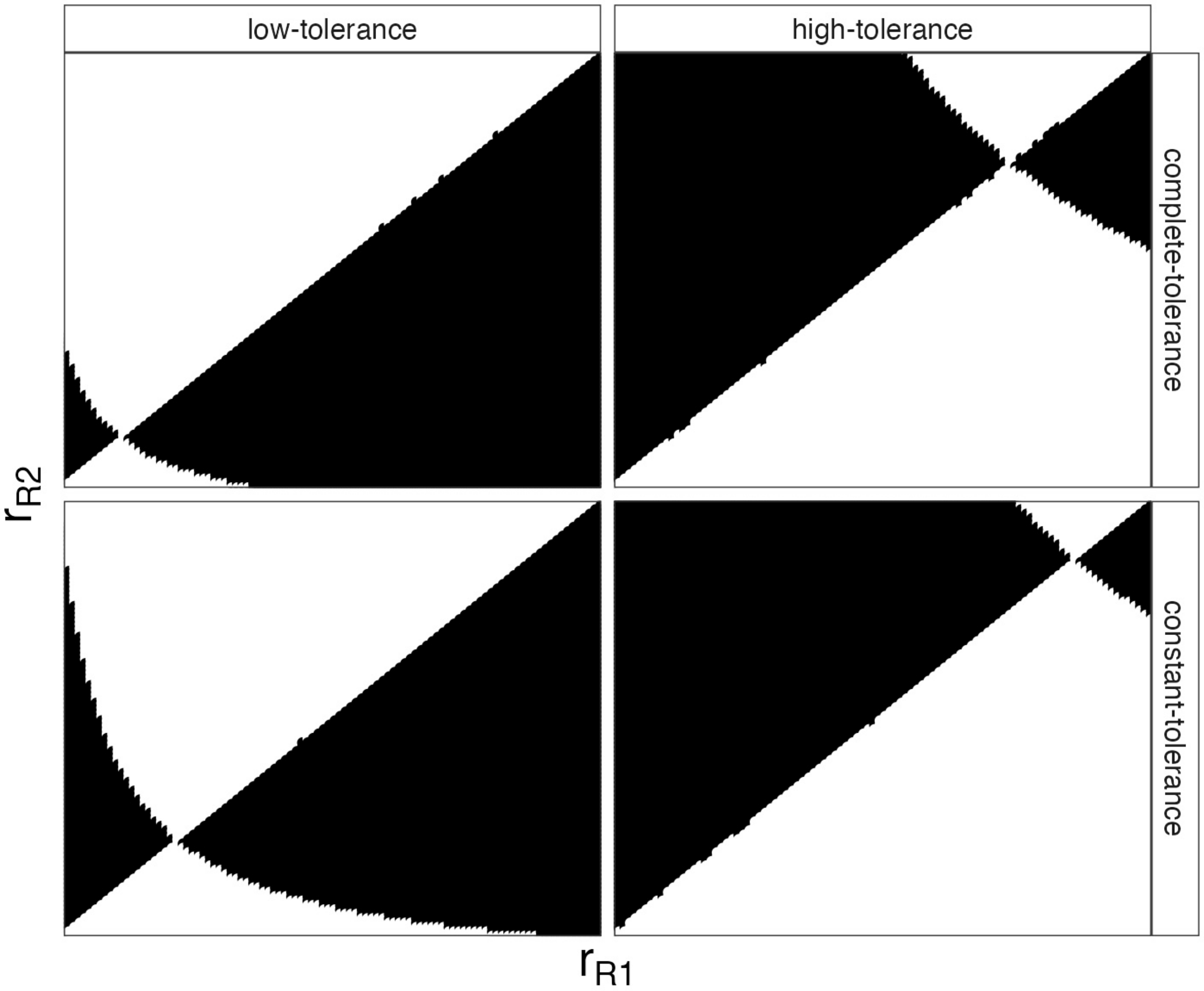
low-tolerance
high-tolerance
complete-tolerance
constant-tolerance
$r_{R2}$
$r_{R1}$

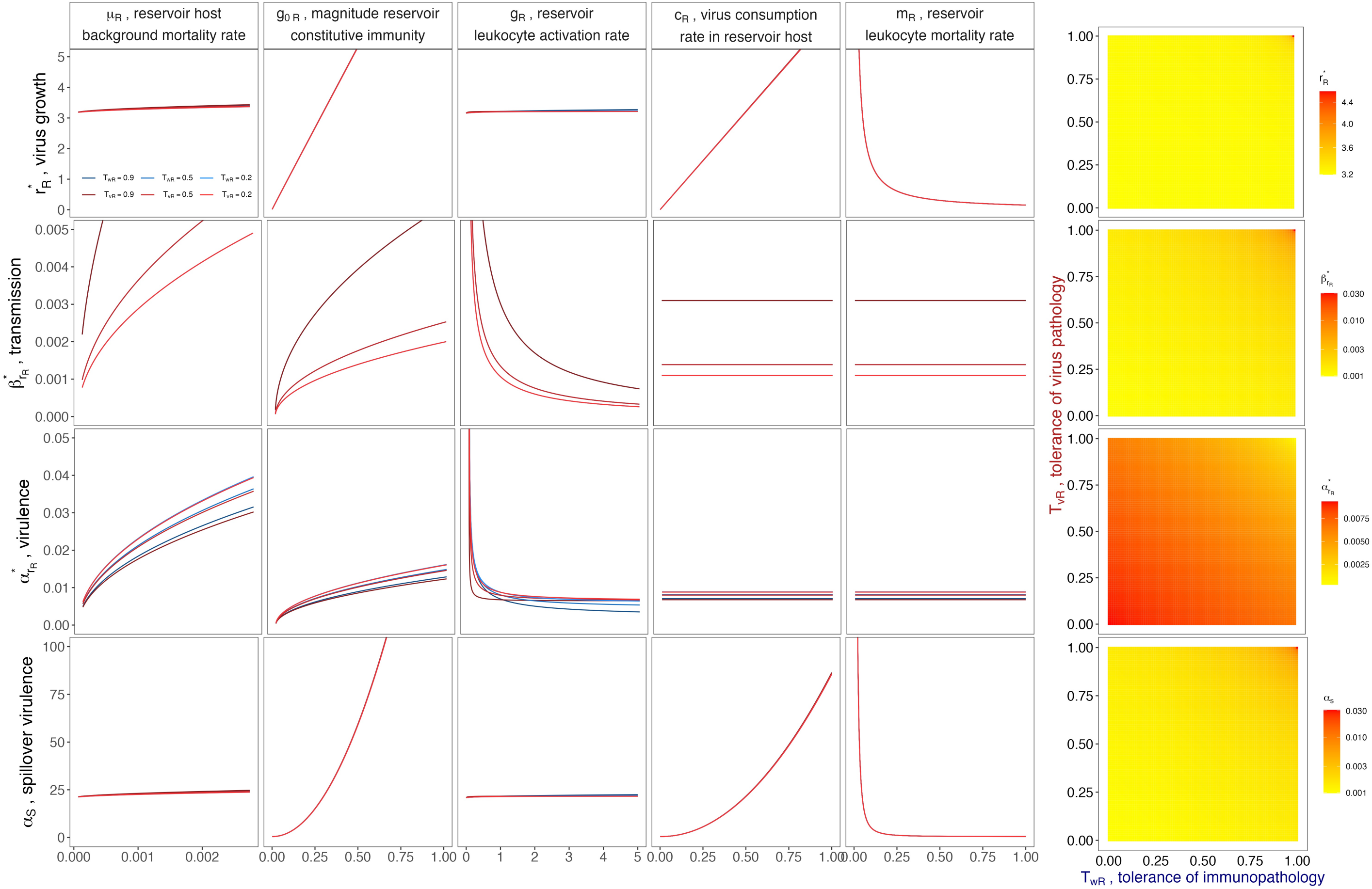

μR , reservoir host background mortality rate
g0 R , magnitude reservoir constitutive immunity
gR , reservoir leukocyte activation rate
cR , virus consumption rate in reservoir host
mR , reservoir leukocyte mortality rate
r*R , virus growth
β*rR , transmission
α*rR , virulence
αS , spillover virulence
TwR = 0.9
TwR = 0.5
TwR = 0.2
TvR = 0.9
TvR = 0.5
TvR = 0.2
TvR , tolerance of virus pathology
TwR , tolerance of immunopathology

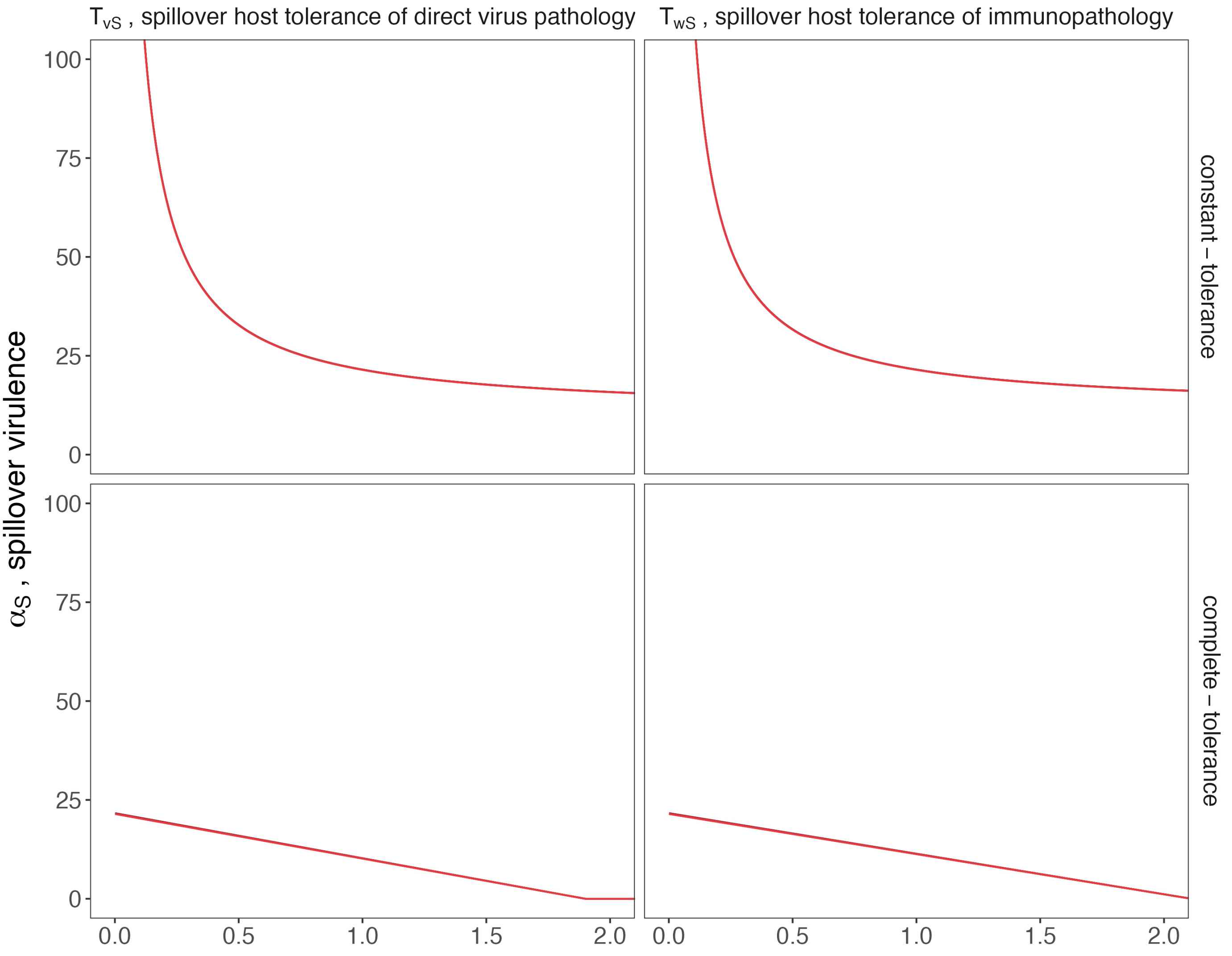

$T_{vS}$ , spillover host tolerance of direct virus pathology
$T_{wS}$ , spillover host tolerance of immunopathology
$\alpha_S$ , spillover virulence
constant – tolerance
complete – tolerance
100
75
50
25
0
0.0
0.5
1.0
1.5
2.0

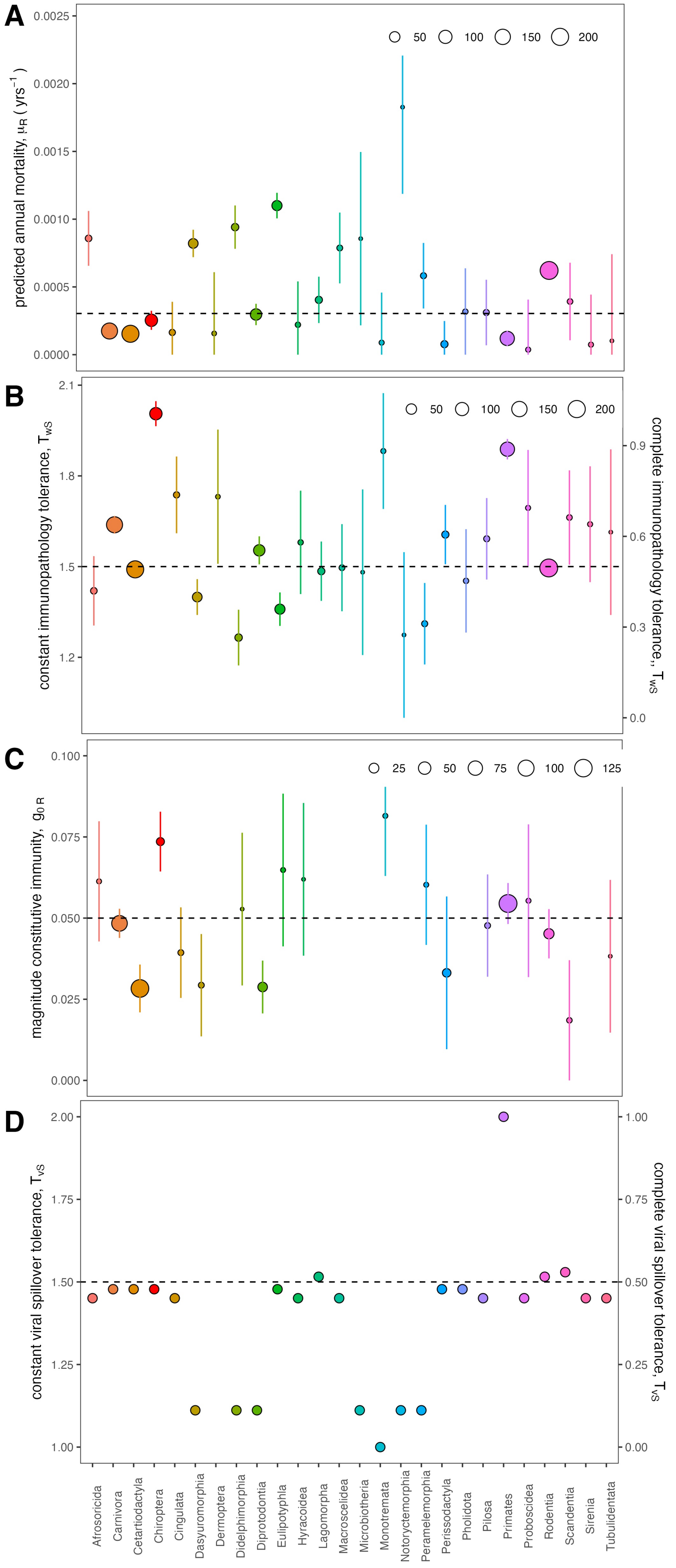

A
B
C
D
predicted annual mortality, μR ( yrs⁻¹ )
constant immunopathology tolerance, TWS
complete immunopathology tolerance, TWS
magnitude constitutive immunity, g0 R
constant viral spillover tolerance, TVS
complete viral spillover tolerance, TVS
50
100
150
200
25
75
125
Afrosoricida
Carnivora
Cetartiodactyla
Chiroptera
Cingulata
Dasyuromorphia
Dermoptera
Didelphimorphia
Diprotodontia
Eulipotyphla
Hyracoidea
Lagomorpha
Macroscelidea
Microbiotheria
Monotremata
Notoryctemorphia
Peramelemorphia
Perissodactyla
Pholidota
Pilosa
Primates
Proboscidea
Rodentia
Scandentia
Sirenia
Tubulidentata

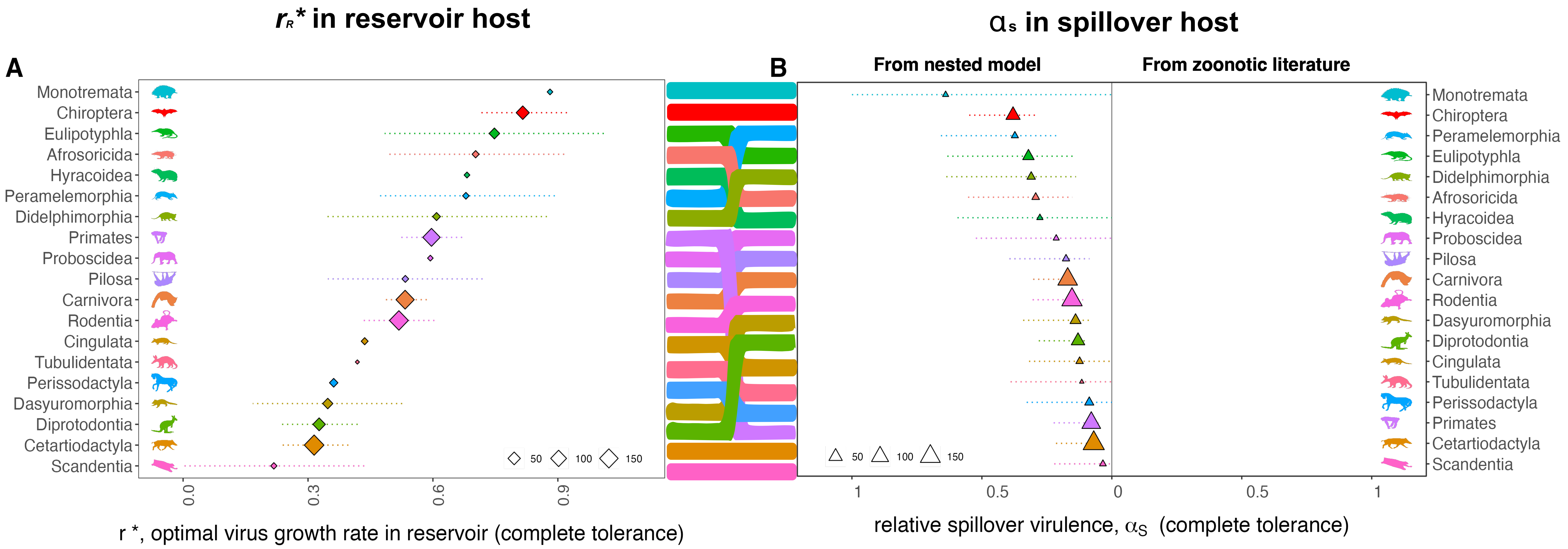

rR* in reservoir host
αs in spillover host
A
B
From nested model
From zoonotic literature
Monotremata
Chiroptera
Eulipotyphla
Afrosoricida
Hyracoidea
Peramelemorphia
Didelphimorphia
Primates
Proboscidea
Pilosa
Carnivora
Rodentia
Cingulata
Tubulidentata
Perissodactyla
Dasyuromorphia
Diprotodontia
Cetartiodactyla
Scandentia
50
100
150
0.0
0.3
0.6
0.9
r *, optimal virus growth rate in reservoir (complete tolerance)
1
0.5
0
0.5
1
relative spillover virulence, αS (complete tolerance)

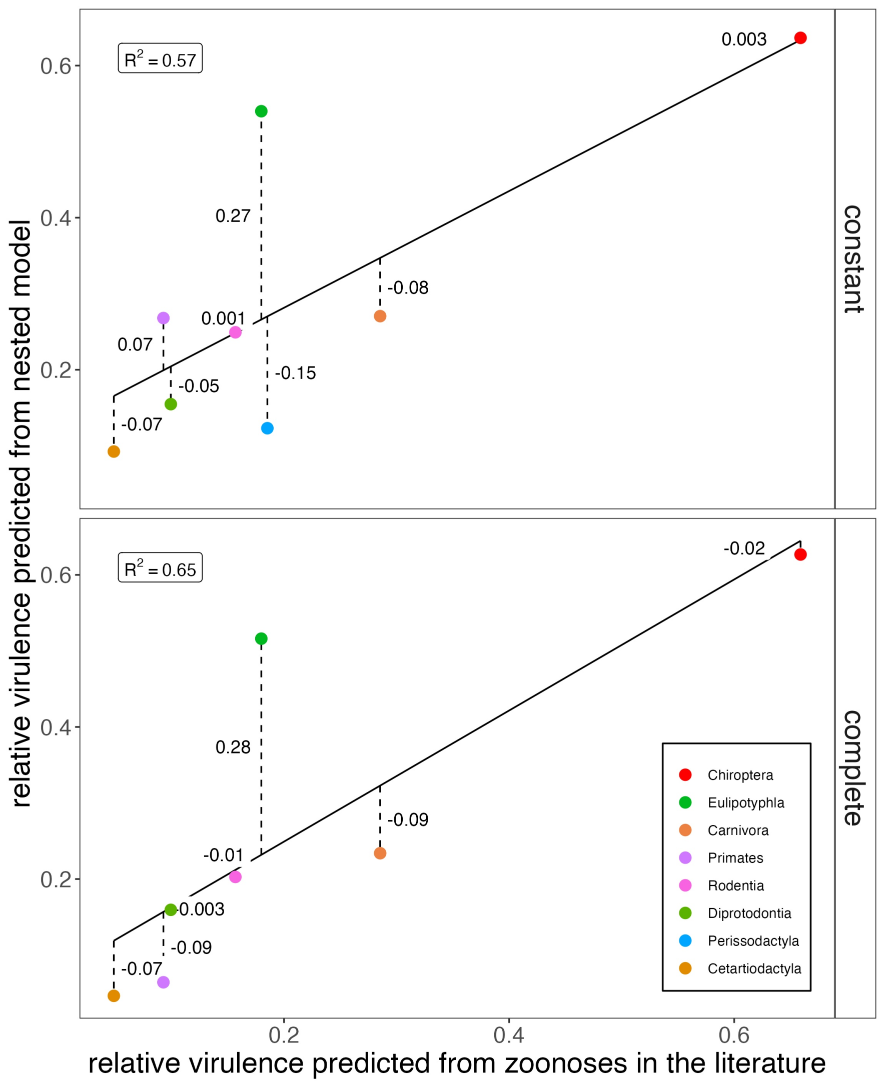

relative virulence predicted from nested model
relative virulence predicted from zoonoses in the literature
constant
complete
$R^2 = 0.57$
$R^2 = 0.65$
0.6
0.4
0.2
0.003
0.27
-0.08
0.001
0.07
-0.05
-0.15
-0.07
-0.02
0.28
-0.09
-0.01
0.003
-0.09
-0.07
Chiroptera
Eulipotyphla
Carnivora
Primates
Rodentia
Diprotodontia
Perissodactyla
Cetartiodactyla

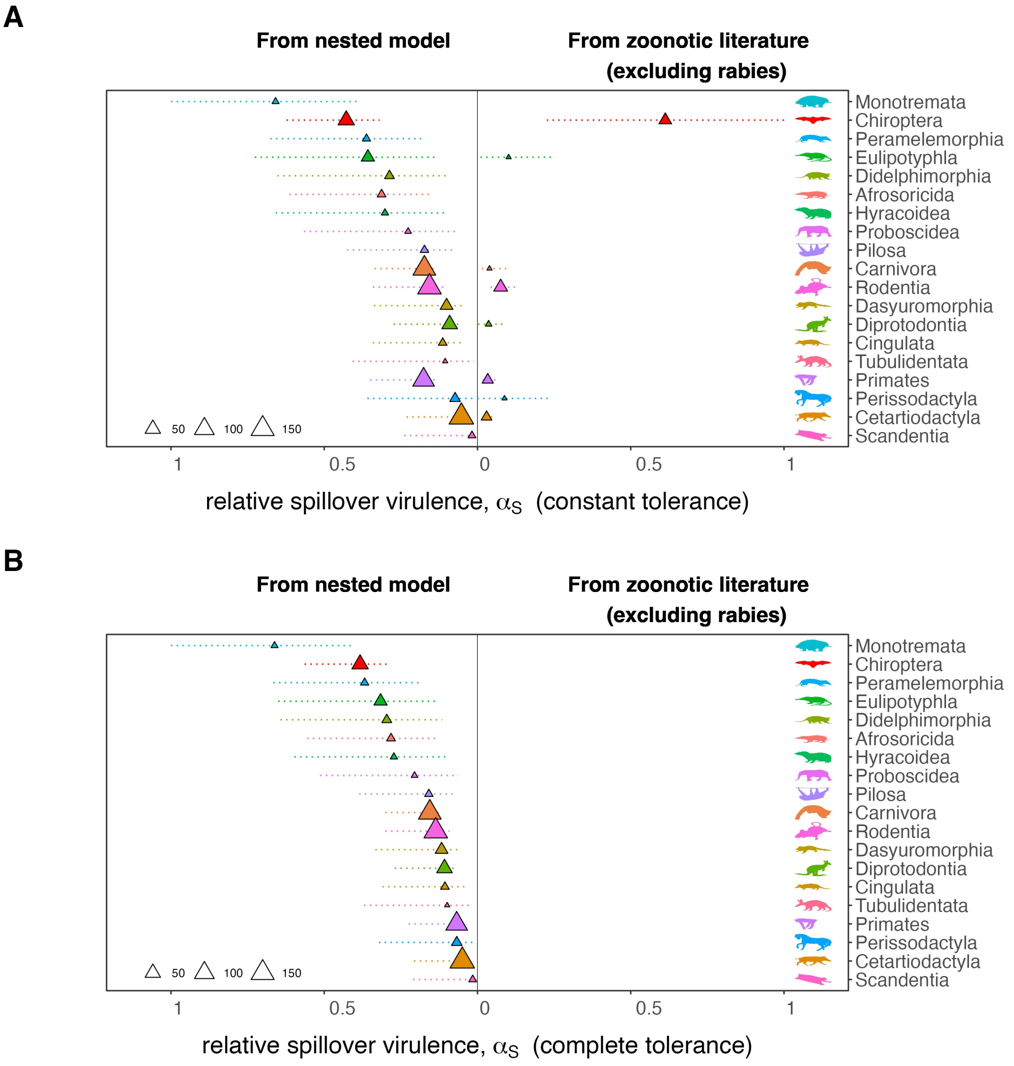

A
From nested model
From zoonotic literature
(excluding rabies)
Monotremata
Chiroptera
Peramelemorphia
Eulipotyphla
Didelphimorphia
Afrosoricida
Hyracoidea
Proboscidea
Pilosa
Carnivora
Rodentia
Dasyuromorphia
Diprotodontia
Cingulata
Tubulidentata
Primates
Perissodactyla
Cetartiodactyla
Scandentia
50
100
150
1
0.5
0
0.5
1
relative spillover virulence, $\alpha_S$ (constant tolerance)
B
From nested model
From zoonotic literature
(excluding rabies)
Monotremata
Chiroptera
Peramelemorphia
Eulipotyphla
Didelphimorphia
Afrosoricida
Hyracoidea
Proboscidea
Pilosa
Carnivora
Rodentia
Dasyuromorphia
Diprotodontia
Cingulata
Tubulidentata
Primates
Perissodactyla
Cetartiodactyla
Scandentia
50
100
150
1
0.5
0
0.5
1
relative spillover virulence, $\alpha_S$ (complete tolerance)

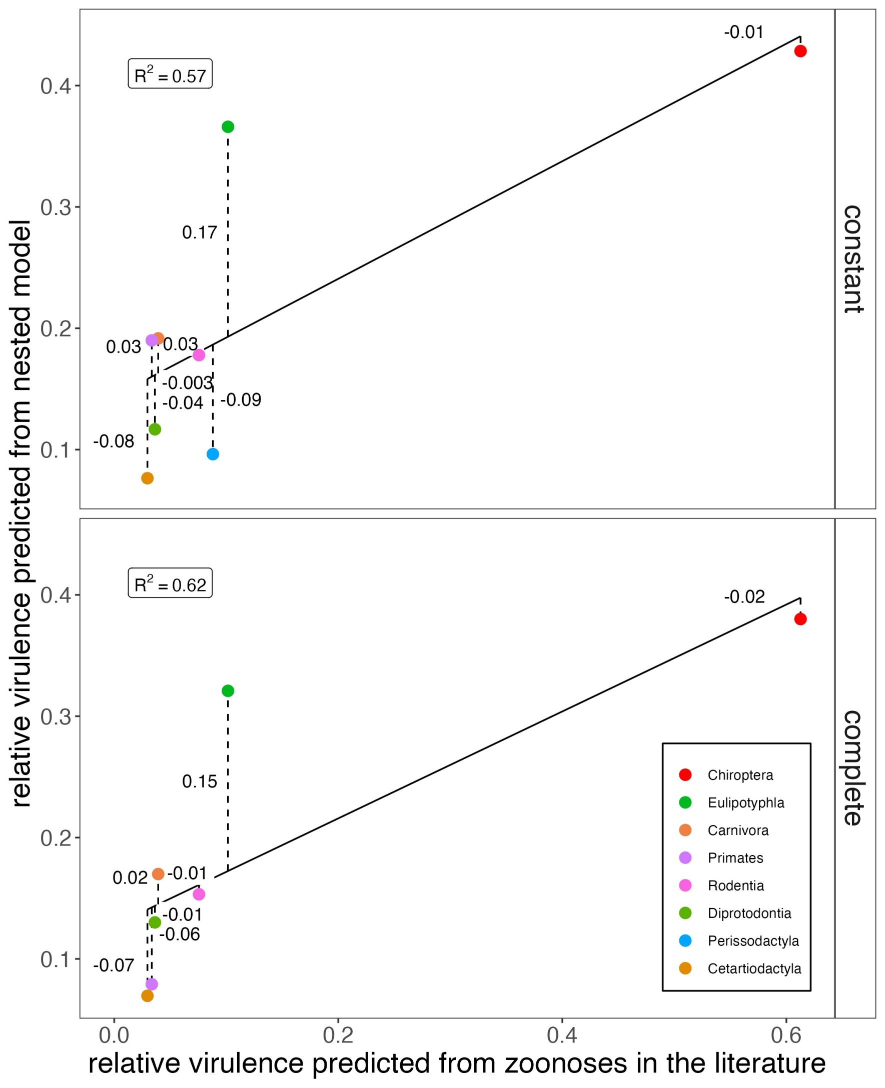

relative virulence predicted from nested model
relative virulence predicted from zoonoses in the literature
constant
complete
$R^2 = 0.57$
$R^2 = 0.62$
-0.01
0.17
0.03
0.03
-0.003
-0.04
-0.09
-0.08
-0.02
0.15
0.02
-0.01
-0.01
-0.06
-0.07
0.1
0.2
0.3
0.4
0.0
0.2
0.4
0.6
Chiroptera
Eulipotyphla
Carnivora
Primates
Rodentia
Diprotodontia
Perissodactyla
Cetartiodactyla

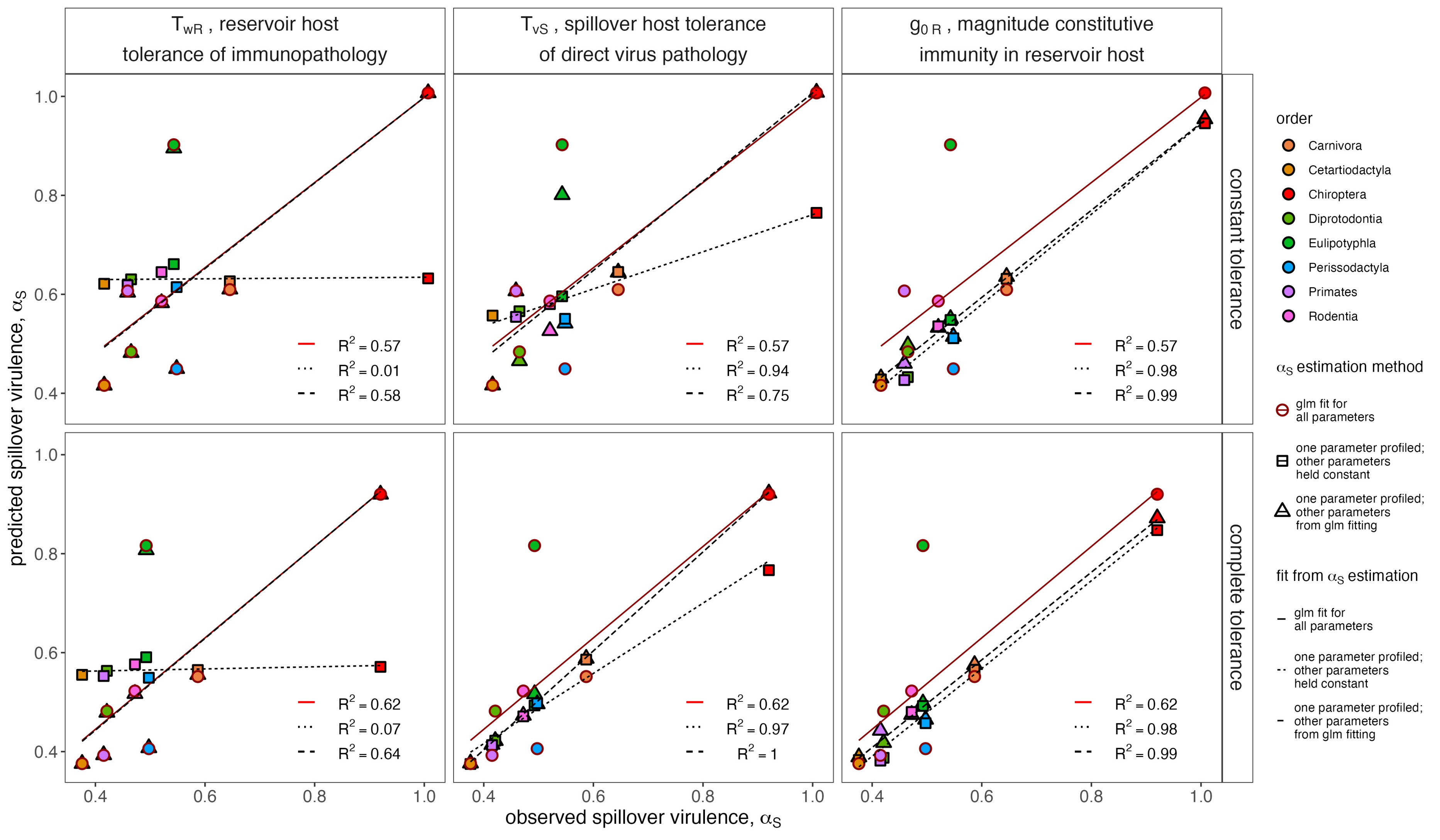

$T_{wR}$ , reservoir host tolerance of immunopathology
$T_{vS}$ , spillover host tolerance of direct virus pathology
$g_{0R}$ , magnitude constitutive immunity in reservoir host
constant tolerance
complete tolerance
predicted spillover virulence, $\alpha_S$
observed spillover virulence, $\alpha_S$
$R^2 = 0.57$
$R^2 = 0.01$
$R^2 = 0.58$
$R^2 = 0.57$
$R^2 = 0.94$
$R^2 = 0.75$
$R^2 = 0.57$
$R^2 = 0.98$
$R^2 = 0.99$
$R^2 = 0.62$
$R^2 = 0.07$
$R^2 = 0.64$
$R^2 = 0.62$
$R^2 = 0.97$
$R^2 = 1$
$R^2 = 0.62$
$R^2 = 0.98$
$R^2 = 0.99$
order
Carnivora
Cetartiodactyla
Chiroptera
Diprotodontia
Eulipotyphla
Perissodactyla
Primates
Rodentia
$\alpha_S$ estimation method
glm fit for all parameters
one parameter profiled; other parameters held constant
one parameter profiled; other parameters from glm fitting
fit from $\alpha_S$ estimation
glm fit for all parameters
one parameter profiled; other parameters held constant
one parameter profiled; other parameters from glm fitting
